# Cell-cycle inhibition preserves robust development but rebalances lineages in mouse gastruloids

**DOI:** 10.64898/2026.01.08.698406

**Authors:** Maxine Leonardi, Yves Paychère, Felix Naef

**Affiliations:** Laboratory of Computational and Systems Biology, Institute of Bioengineering, Faculty of Life Sciences, É cole Polytechnique Fédérale de Lausanne (EPFL), Lausanne, Switzerland

**Keywords:** Cell cycle, Gastruloid, Gastrulation, Development, Cell differentiation, Cell proliferation, Stem cells, scRNAseq, modelling

## Abstract

Cell differentiation and proliferation are fundamental to the development of multicellular organisms. While studies in various non-mammalian species show that development can proceed despite disrupted cell cycle progression, the extent to which normal cell cycle dynamics are required in mammals remains unclear. Using mouse gastruloids, we examined the effects of cell cycle inhibition on development. Despite near-complete growth arrest, gastruloids still underwent symmetry breaking, elongation, and germ layer specification, indicating that core differentiation programs are robust to growth inhibition. However, microscopy and single-cell transcriptomics revealed consistent alterations in cell type proportions, including delayed differentiation and reduced mesodermal populations. To investigate the origin of these changes, we used the differential kinetics of unspliced and spliced cycling transcripts to compare proliferation rates between cell types. While differences in proliferation partly explained the imbalance, our analysis showed that cell cycle perturbations also modulate lineage-specific differentiation timing and efficiency, highlighting a regulatory role of cell cycle dynamics in mammalian development.

## INTRODUCTION

The cell cycle is a fundamental process that governs the growth, development, homeostasis, and regeneration of all living organisms, and its dysregulation is a hallmark of many diseases^1,2^. Beyond the primary role of the cell cycle in growth control, its direct involvement in regulating pluripotency and cell fate commitment has become increasingly clear. One of the most consistent observations is that higher potential for pluripotency coincides with a faster cell cycle, often marked by a shortened G1 phase^3^. At the molecular level, core pluripotency factors directly promote the cell cycle by acting upstream of key cell cycle regulators^4–12^. Fast cell cycles are not only a hallmark of pluripotent cells but also actively sustain this state. Indeed, cell cycle regulators are, in turn, direct regulators of pluripotency factors^13–17^. Therefore, inhibition or knockout of many CDKs or cyclins leads to spontaneous differentiation^18–23^, while their overexpression can delay lineage commitment^22,24^.

Cell cycle control remains essential throughout the transition from pluripotency to differentiation. As cells differentiate, the cell cycle duration extends with the G1 phase becoming longer, providing a window of opportunity to integrate external differentiation signals, partly through chromatin restructuring^25,26^. This extended G1 phase along with the proteins that regulate it have been shown to play a crucial role in the commitment towards multiple lineages, such as mesoderm^27,28^, neural^29–32^, and endothelial tissues^33^. Moreover, the timing of differentiation cues within G1 phase also influences cell fate: exposure to such cues in early G1 promotes mesendodermal fate, whereas cues in late G1 favor neuroectoderm, partly via CDK-dependent regulation of cell signaling pathways^34,35^. Beyond the cell cycle regulation of individual cell fates, early development in many species, such as *Zebrafish*, *C. elegans* and *Xenopus*, is characterized by synchronized and spatially coordinated cycle events^36–38^. Similarly, a functional role of the spatial organisation of cell cycle phases has been identified in mouse development, particularly during somitogenesis^39,40^.

These works highlight a clear interplay between the cell cycle and cell fate decisions. Maintaining the balance between pluripotency and differentiation is essential during early embryogenesis. Therefore, altering cell cycle progression should potentially derail this balance and impair normal development. However, evidence across model organisms suggests that early development is surprisingly robust to cell cycle perturbations^41^. Indeed, in several systems, early development proceeds despite arrested proliferation: in *C. elegans*, cell cycle mutants that abolish mitosis can still initiate transcriptional programs of differentiation on time^42^. Similarly, in *Xenopus*, blocking DNA replication does not prevent morphogenetic events like neurulation^43–45^. In *Zebrafish*, embryos can still undergo axis elongation and form blood, muscle, and a beating heart even when cell division is blocked from gastrulation onward. However, this arrest in proliferation leads to defects in morphogenesis, cell migration, and tissue organization^46,47^. A recent single-cell RNA-seq study has further confirmed that differentiation of all major embryonic tissues can occur in the absence of cell division^48^. However, the proportions of the resulting cell types were altered significantly, with differences of up to tenfold, and some lineages showed a delay in differentiation, indicating that while proliferation is not strictly required for lineage specification, it does influence the pace and balance of developmental processes.

In mouse, most cyclins and CDKs are dispensable for early development, with knockouts causing only limited or tissue-specific defects^49–51^. However, these mild phenotypes likely arise from functional redundancy among cell cycle regulators, which maintains near-normal cell cycle dynamics during embryogenesis, rather than from an inherent robustness of development to cell cycle perturbations. Effective perturbations of the cell cycle has also been investigated in mouse embryos through transient chemical arrest of proliferation, which significantly reduced cell numbers, but still allowed embryos to undergo morphogenesis, although the long-term consequences include impaired growth and fertility^52^. In other studies, altering cell cycle dynamics in the mouse embryo has been shown to cause defects in specific tissues, notably in the neuronal lineage^53,54^, or in later developmental processes such as somitogenesis, where cell cycle arrest leads to abnormalities in somite length^40^. In humans, mutations in CDKs or their inhibitors have been linked to developmental disorders such as microcephaly or gigantism^55–57^. Moreover, in vitro studies using human embryoid bodies indicate that altering cell cycle regulators can shift lineage allocation^58^. Together, these findings illustrate a complex, context-dependent relationship between proliferation and mammalian development, and highlight that the overall influence of the cell cycle on cell fate specification remains poorly understood, particularly in mammalian development.

Here, we use mouse *gastruloids*, an in-vitro model of gastrulation^59–65^, to investigate the consequences of cell cycle inhibition on mammalian development. Combining microscopy and scRNA-seq, we show that despite near-complete proliferation arrest after drug treatment, gastruloids preserved key developmental features such as symmetry breaking and elongation, and generated cell types from all three germ layers. However, cell type proportions were quantitatively altered in growth-arrested gastruloids consistently across all data modalities, with delayed differentiation in specific lineages and a rebalancing of the distributions of cell differentiation trajectories. To investigate the underlying cause of these differences, we compare unspliced and spliced RNA profiles to map cell cycle rates along the different differentiation lineages. This demonstrated that differences in proliferation rates partially accounted for the changes observed in growth-arrested gastruloids, but were insufficient to fully explain them, arguing that the cell cycle also likely modulates the efficiency of differentiation through another mechanism. Together, these findings suggest that although the cell cycle is not required for the developmental processes at play during gastrulation, it likely contributes to lineage balance and timing within mouse gastruloids.

## RESULTS

### 0.1 Cell Cycle-inhibited gastruloids break symmetry, elongate, and recapitulate patterned marker gene expression but show altered cell type proportions

To investigate how modulating cell cycle kinetics influences early mammalian development, we generated gastruloids from mouse embryonic stem cells using standard protocols^59^ and treated with pharmacological inhibitors targeting key regulators of proliferation, namely CDK4/6, CDK2, CDK1 and mTOR from either 48h or 72h after aggregation (A.A.) onward (**Figure 1A**). These conditions and doses were titrated to slow down the cell cycle without inducing gastruloid death. In addition, a Wnt pathway inhibitor was used to probe abnormal development, as it is expected to impair gastruloid development by acting directly upstream of T, a major regulator of mesendo-dermal fated cells. Gastruloid growth and morphology were monitored by brightfield imaging every 24h (**Figure 1B**, **Supp. Figure S1A**), and samples were collected at 120h A.A. for subsequent immunofluorescence staining (IF) (**Figure 1C**).

**Figure 1.**
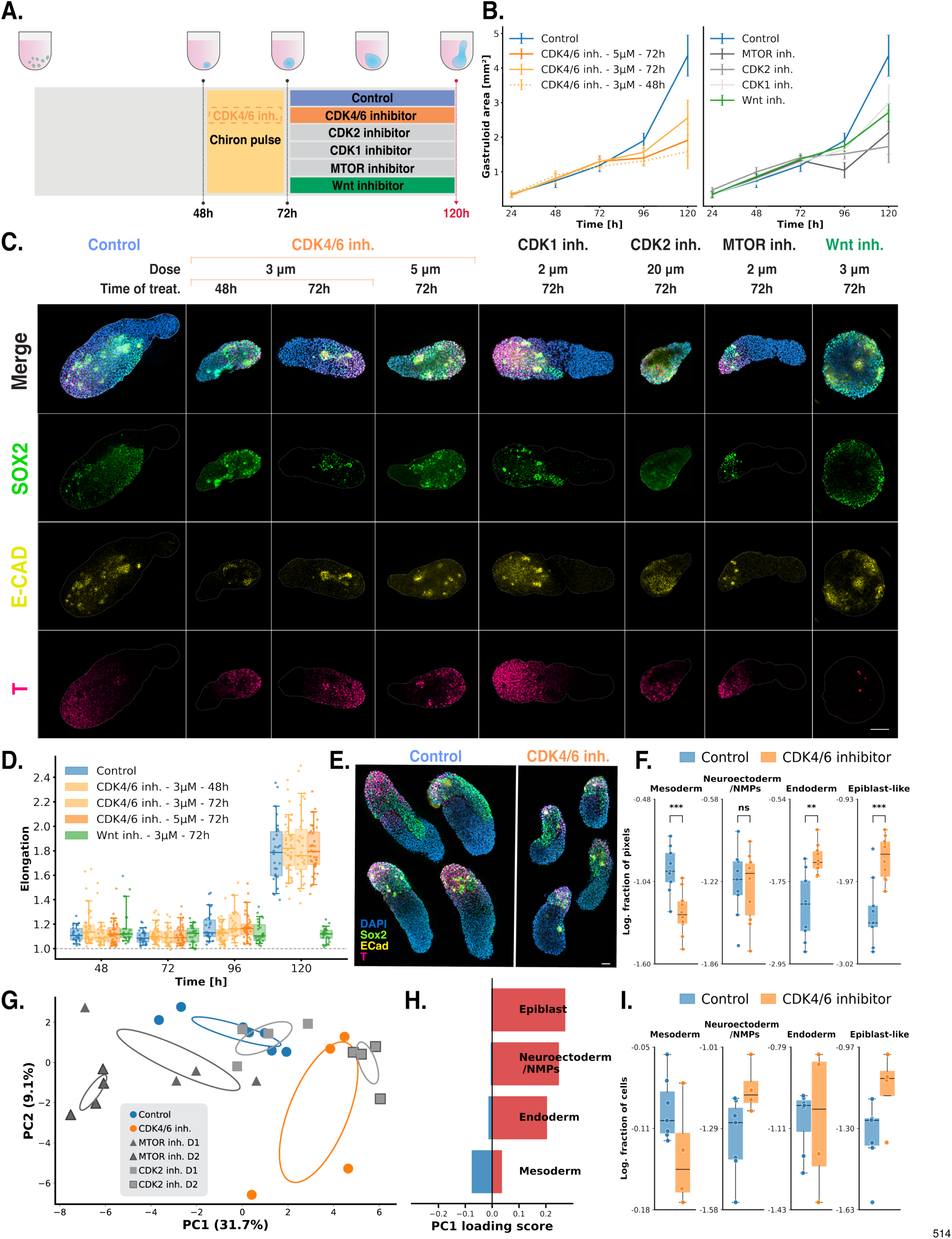
Cell Cycle inhibited gastruloids break symmetry, elongate, and recapitulate patterned marker gene expression, but show altered cell type proportions. (A) Experimental design: Gastruloids were generated by aggregating 300 mouse ES cells and treating them with Chiron from 48h to 72h. From 48h or 72h onward, they were treated with either DMSO (Control), CDK4/6 inhibitor (Palbociclib), CDK2 inhibitor (NU6102), CDK1 inhibitor (RO3306), MTOR inhibitor (Torin1) or Wnt inhibitor (Porcupine). Gastruloids were imaged with the Operetta microscope every 24h, and collected at 120h for subsequent immunofluorescence stainings. (B) Operetta-based quantification of gastruloid area progression reveals substantially reduced growth under CDK4/6 (left) and other cell cycle and Wnt inhibitor treatments (right). *n >*= 20 for all conditions. Error bars represent the standard deviation (SD). (C) IF of gastruloids in control and drugged conditions. Stainings for T, SOX2, and E-CAD show that cell cycle inhibition does not prevent cell type marker expression and patterning, despite strong size reduction. Scale = 100 µm. (D) Quantification of elongation reveals strong impairment under Wnt inhibition, but no effect with CDK4/6 inhibition. *n >*= 20 for all conditions. (E) Further representative staining of T, SOX2, and E-CAD in control and CDK4/6-inhibited gastruloids (5µM, from 72 h), the conditions selected for subsequent quantification. Scale = 50 µm. (F) Comparison of cell type composition from IF stainings in control and CDK4/6-inhibited gastruloids. Quantification shows a significant reduction of mesodermal cells and an increase of early tissue populations retaining epiblast-like pluripotency. *n*_control_ =9, *n*_palbociclib_ = 10. Statis-tical significance was assessed using a two-sided Welch’s t-test on log-transformed data, with *p <* 0.05 (*), *p <* 0.01 (**), and *p <* 0.001 (***) indicating significant differences. (G) PCA of cell type marker genes from bulk sequencing of single gastruloids across control and inhibitor treatments. Samples cluster by condition along PC1, reflecting distinct cell type marker expression profiles. *n*_control_ =7, *n*_others_ = 4. (H) PC1 gene loading scores of cell type marker genes. The score (sum of squared loading) reflects the contribution of marker genes from each cell type to PC1. Epiblast, endodermal, and neuroectodermal markers load positively on PC1. Mesodermal markers load negatively on PC1. (I) Cell type proportions from bulk RNA sequencing after cell type deconvolution with Scaden^68^ reveals trends consistent with immunostaining quantifications (E).

Across all cell cycle inhibition conditions, gastruloid growth was strongly reduced after treatment compared to controls, with consistent results across replicates (**Figure 1B**, **Supp. Figure S1B**). In particular, the higher doses of CDK4/6 inhibitor (5µM) produced gastruloids approximately 3 times smaller in area. In this condition, the gastruloid area continued to increase slightly over time, even under drug treatment. However, this likely reflects their elongated shape at 120 h, which results in a larger projected area for the same volume rather than an increase in cell number. Consistent with this interpretation, quantification of cell counts over time showed no net growth after treatment, indicating a balance between proliferation and apoptosis in the presence of the drug (**Supp. Figure S2B**).

Interestingly, these growth-arrested gastruloids still underwent symmetry breaking despite their reduced size. This process, defined as the emergence of a molecular or morphological asymmetry from an initially homogeneous cell population, was evident from the formation of an axis and the polarised expression of marker genes in all cell cycle inhibited gastruloids. Elongation, quantified as the ratio between the gastruloid’s major and minor axes (**Supp. Figure S1C**), showed no significant difference between control and CDK4/6-inhibited gastruloids, whereas Wnt inhibition completely abolished elongation, with gastruloids that stayed spherical until 120h (**Figure 1D**). These results indicate that normal cell cycle dynamics are not required for symmetry breaking or axis formation in gastruloids.

Moreover, cell type markers of all the germ layers (combination of T, SOX2, E-CAD) were still expressed in all cell cycle inhibited gastruloids at 120h, forming clusters and patterns similar to those observed in control gastruloids (**Figure 1C**). In contrast, Wnt-inhibited gastruloids almost completely failed to express the mesodermal marker T, nor did they show regions of E-CAD expression without SOX2, an endodermal marker, indicating the expected disrupted mesendoderm formation. Together, these findings suggest that cell cycle inhibition does not markedly impair either the emergence or the patterning of distinct germ layer cell types in gastruloids.

In the following analyses, one representative cell cycle inhibition condition (CDK4/6 inhibition at 5µM from 72h) was selected for further quantitative assessment of cell type proportions.

In this condition, the effects of the inhibitor on cell cycle dynamics were characterised using several complementary approaches. This included Fucci(CA) imaging and flow cytometry analyses performed throughout gastruloid development (**Supp. Figure S2A-E**). In this reporter, CDT1 expression marks cells in the G1 phase, GEMININ expression marks cells in the S phase, and co-expression of both indicates that cells are in the G2/M phase. Quantification from microscopy images and flow cytometry data both showed a time-dependent accumulation of G1-phase cells in CDK4/6-inhibited gastruloids compared to controls (**Supp. Figure S2C,E**). This phenotype was further confirmed by DNA content profiles derived from DAPI fluorescence in the flow cytometry experiments (**Supp. Figure S2D,H**). These findings are consistent with CDK4/6 acting as a G1/S checkpoint regulator. Although an enrichment of G1 cells is observed, other cell cycle phases are still represented in the drugged gastruloids, indicating that cell cycle is slowed down rather than completely blocked in the G1 phase.

To assess whether other phases of the cell cycle were also affected by cell cycle inhibition, proliferation assays were performed by pulsing 120h gastruloids with 5-ethynyl-2’-deoxyuridine (EdU) for 2h. Imaging and flow cytometry analyses both revealed a strong decrease in EdU incorporation in the inhibited condition (**Supp. Figure S2F,H,I,J**). Moreover, the EdU-positive cells that remained exhibited lower fluorescence intensity than controls (**Supp. Figure S2J**), suggesting that not only fewer cells entered S phase, but also that those cells replicated their genomes more slowly. Together, these results demonstrate that CDK4/6 inhibition induces a globally slower cell cycle progression, with a particular cell accumulation in the G1 phase. Moreover programmed cell death, which has been showed to be a main driver of cell competition in gastruloid^66^, was variable under the CDK4/6 inhibition between gastruloids, but tended to be slightly higher overall than in controls, as shown by activated caspase staining (**Supp. Figure S2G,K**).

Having established the effects of CDK4/6 inhibition on cell cycle progression, we next investigated whether this perturbation also impacted the cell type composition of gastruloids using cell marker immunofluorescence staining (**Figure 1E**). Automated quantification of immunostainings using a pixel-based classification (**Supp. Figure S1D, E**) revealed significant shifts in lineage proportions. Specifically, CDK4/6 inhibition reduced the fraction of T-positive mesodermal cells while increasing both E-CAD-positive populations and SOX2 and E-CAD-positive populations characteristic of endoderm and epiblast-like early tissues, respectively (**Figure 1F**).

To complement these observations, we next asked whether these lineage shifts were also reflected at the transcriptional level. Therefore, we performed bulk RNA sequencing of individual gastruloids at 120h, in control conditions and in conditions where the cell cycle is inhibited with multiple drugs (CDK4/6 inhibitor, CDK2 inhibitor., mTOR inhibitor). In the CDK4/6 inhibited condition, differential expression analysis confirmed transcriptional changes related to cell cycle regulation, including the up-regulation of D-type cyclins and down-regulation of proliferation markers such as *Mki67* and *Pcna* transcripts (**Supp. Figure S2L-M**, **Supp. Table 1**). The increased expression of D-cyclins is consistent with the CDK4/6 inhibitor (Palbociclib) signature observed previously^67^, and likely reflects a compensatory response following inhibition of their regulatory partners. Principal component analysis (PCA) of cell type marker genes separated samples by condition, indicating that marker gene expression profiles differ between treatments (**Figure 1G**). Interestingly, CDK4/6 and CDK2 inhibition, both acting on the G1/S checkpoint, exhibited opposite trends to mTOR inhibition relative to the control, suggesting that these pathways may have opposing effects on cell type composition. The gene loadings of principal component one (PC1) indicated that mesodermal markers contributed negatively to this axis, whereas epiblast, neuroectodermal/NMPs, and endodermal markers loaded it positively (**Figure 1H**). Thus, the positive shift of CDK4/6-inhibited gastruloids relative to controls is consistent with a decrease in mesodermal and an increase in endodermal and epiblast-like transcriptional signatures. Moreover, cell type deconvolution using Scaden^68^ corroborated this, confirming a decrease in mesodermal fractions and enrichment of epiblast-like populations consistent with IF quantifications (**Figure 1I**).

**Table 1:**
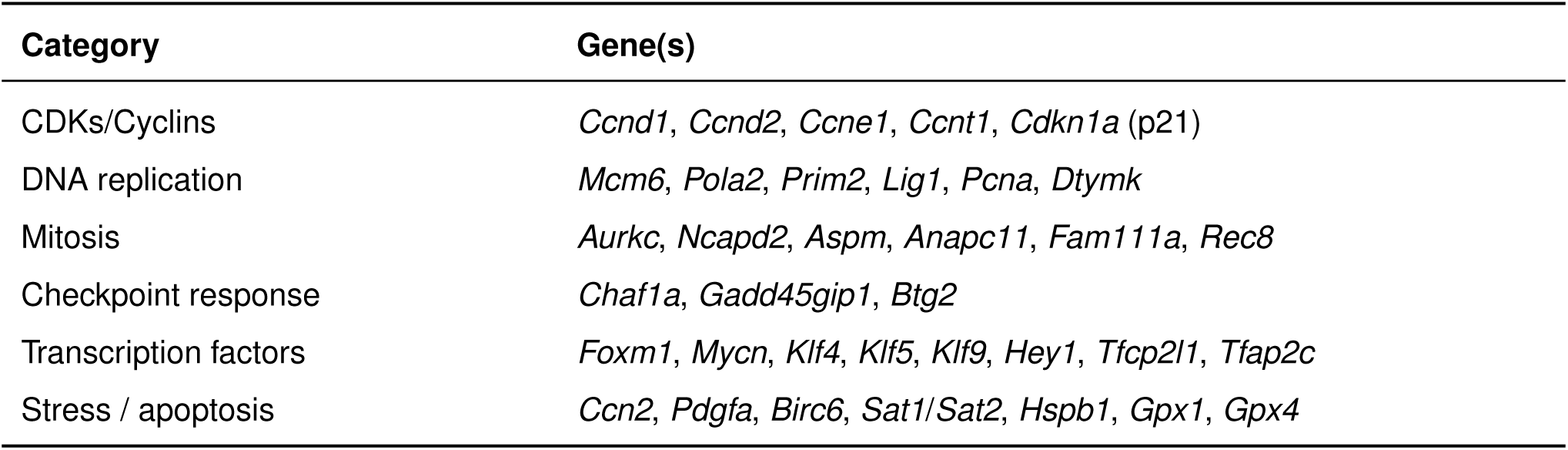
Differentially expressed genes from bulk RNA-seq comparing control and CDK4/6-inhibited conditions indicate disruption of cell-cycle regulation.

| Category | Gene(s) |
| --- | --- |
| CDKs/Cyclins | *Ccnd1, Ccnd2, Ccne1, Ccnt1, Cdkn1a* (p21) |
| DNA replication | *Mcm6, Pola2, Prim2, Lig1, Pcna, Dtymk* |
| Mitosis | *Aurkc, Ncapd2, Aspm, Anapc11, Fam111a, Rec8* |
| Checkpoint response | *Chaf1a, Gadd45gip1, Btg2* |
| Transcription factors | *Foxm1, Mycn, Klf4, Klf5, Klf9, Hey1, Tfcp2l1, Tfap2c* |
| Stress / apoptosis | *Ccn2, Pdgfa, Birc6, Sat1/Sat2, Hspb1, Gpx1, Gpx4* |

Taken together, these results show that gastruloid development remains remarkably robust under conditions of cell cycle slow-down. Even when the global gastruloid growth is halted, gastruloids can break symmetry, elongate, and establish patterned germ layer domains. Moreover, expression of the main germ layer markers still emerges, indicating that CDK4/6 inhibition does not completely disrupt differentiation. However, lineage proportions appear altered consistently at both the protein (IF) and mRNA levels, suggesting that while normal cell cycle progression is not essential for the initiation of germ layer formation, it contributes to maintaining balanced lineage specification and may influence the efficiency of fate transitions within developing gastruloids.

### 0.2 scRNA-seq reveals that cell cycle inhibition preserves core developmental trajectories but alters lineage balance and delays differentia-tion in gastruloids

To further examine the consequences of cell cycle inhibition on gastruloid development at the single-cell level and to investigate how the different lineage trajectories unfold over time under growth arrest, we performed single-cell RNA sequencing (scRNA-seq) on pooled gastruloids under three previously described conditions (**Methods**): untreated controls (DMSO), CDK4/6-inhibition (5µM), and Wnt-inhibition. Samples were collected at 0h (mESCs) and then every 12 hours from 48h to 120h, with treatments applied from 72h onward (**Figure 2A**). This time-resolved design enabled us to capture transcriptional changes across thousands of individual cells and reconstruct gastruloid development from early to late stages in both normal and per-turbed conditions.

**Figure 2.**
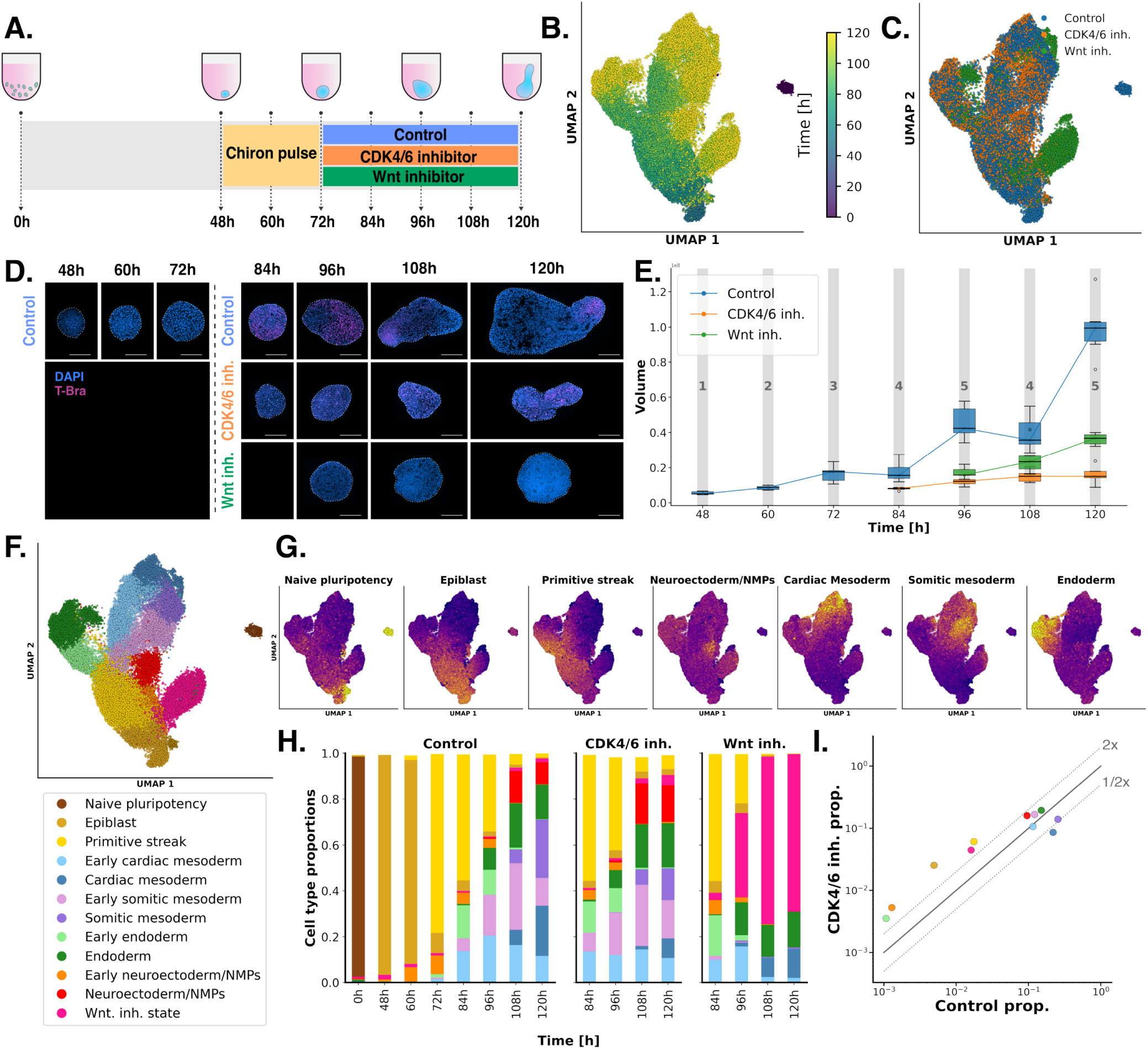
scRNA-seq confirms preserved developmental trajectories but rebalanced cell type proportions in cell cycle-inhibited gastruloids. (A) Experimental design: Gastruloids were generated by aggregating 300 mouse ES cells and treating them with Chiron from 48h to 72h. From 72h onward, they were treated with either DMSO, CDK4/6 inhibitor (Palbociclib), or Wnt inhibitor (Porcupine). For scRNA-seq, gastruloids were pooled and collected at time points indicated by an arrow. (B) UMAP projection of scRNA-seq data showing cell state evolution over time. (C) UMAP projection of scRNA-seq data shows cell state distribution over the different conditions. (D) Microscopy of gastruloids from scRNA-seq batches confirms previously observed phenotypes. (E) Gastruloid volume quantification shows growth arrest under CDK4/6 inhibition, leading to a ninefold smaller size than controls after two days of treatment. (F) UMAP projection of cell type classification of scRNA-seq dataset shows effective differentia-tion into neuroectoderm/NMPs, mesoderm and endoderm trajectories in gastruloids. (G) UMAP projection highlights specific marker gene expression across different cell types. (H) Proportions of the different cell types, stratified by time of collection and conditions reveals similar cell type distribution in Control and CDK4/6 inhibitor conditions, but very distinct popula-tions in Wnt inhibitor condition. (I) Comparison of cell type proportions in Control versus CDK4/6 inhibitor conditions, shown in log scale, reveals a rebalancing of cell populations. Dotted lines indicate a twofold difference.

Imaging of gastruloids from the same batches used for scRNA-seq confirmed the above phenotypes: after treatment onset, CDK4/6-inhibited gastruloids showed no net volume growth, resulting in a final volume approximately 9.5 times smaller than controls, yet they still elongated and polarised normally. Conversely, Wnt-inhibited gastruloids failed to elongate and remained spherical (**Figure 2D,E**, **Supp. Figure S3A**). Moreover, CDK4/6-inhibited cells displayed the expected signatures of cell cycle impairment, including D-type cyclin overexpression and enrichment of genes associated with cell cycle–related GO terms (**Supp. Figure S3C,D**, **Supp. Table 2**).

**Table 2:**
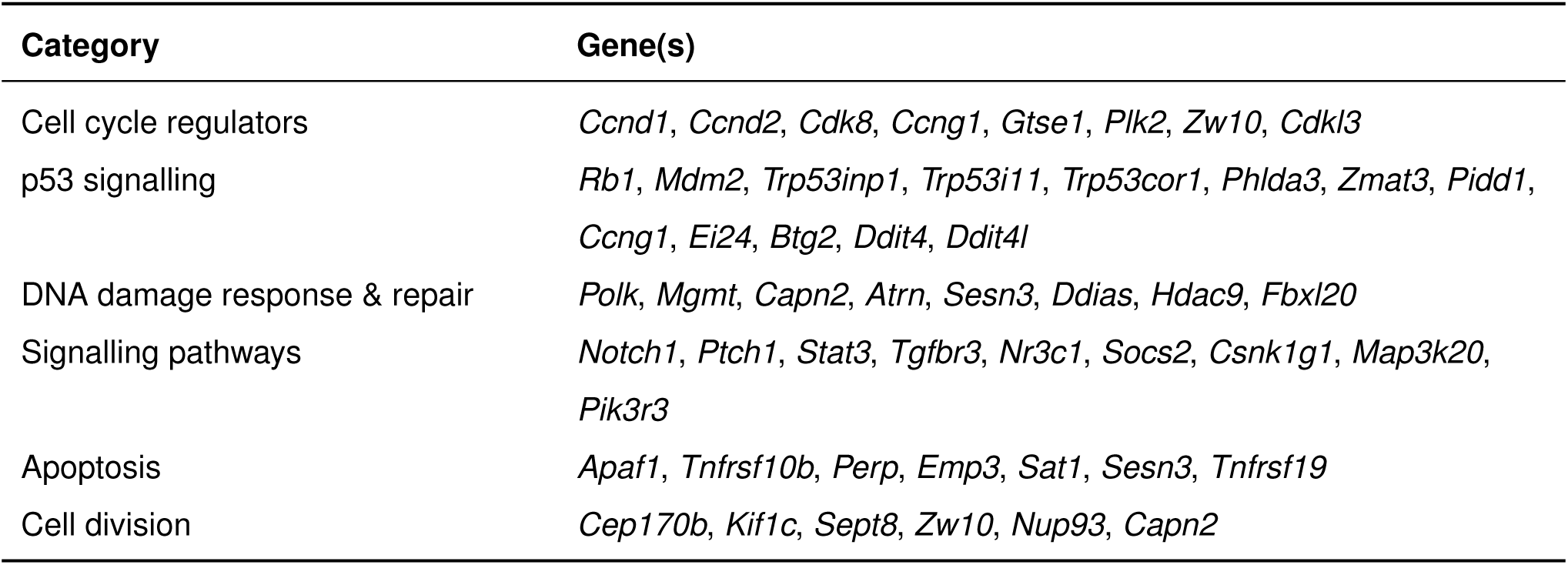
Genes driving classification of control vs CDK4/6-inhibited cells in scRNA-seq data reflect cell-cycle and p53-related responses. Classification was performed using logistic regression for each cell type, and the list represents the most recurrent genes across lineages.

| Category | Gene(s) |
| --- | --- |
| Cell cycle regulators | *Ccnd1, Ccnd2, Cdk8, Ccng1, Gtse1, Plk2, Zw10, Cdkl3* |
| p53 signalling | *Rb1, Mdm2, Trp53inp1, Trp53i11, Trp53cor1, Phlda3, Zmat3, Pidd1, Ccng1, Ei24, Btg2, Ddit4, Ddit4l* |
| DNA damage response & repair | *Polk, Mgmt, Capn2, Atrn, Sesn3, Ddias, Hdac9, Fbxl20* |
| Signalling pathways | *Notch1, Ptch1, Stat3, Tgfbr3, Nr3c1, Socs2, Csnk1g1, Map3k20, Pik3r3* |
| Apoptosis | *Apaf1, Tnfrsf10b, Perp, Emp3, Sat1, Sesn3, Tnfrsf19* |
| Cell division | *Cep170b, Kif1c, Sept8, Zw10, Nup93, Capn2* |

Dimensionality reduction of the scRNA-seq dataset revealed a continuous distribution of cell states consistent with progressive differentiation. All samples, except the 0h time point which formed a distinct cluster corresponding to pluripotent cells, were arranged along a continuum from early to late time points (**Figure 2B**). When comparing conditions, control and CDK4/6-inhibited cells showed largely overlapping distributions over the UMAP space, indicating similar transcriptional trajectories despite growth arrest (**Figure 2C**). In contrast, Wnt-inhibited cells diverged strongly from controls and formed distinct clusters absent in the control condition.

Annotation of cell types using established lineage marker genes revealed three distinct tra-jectories that cells can take, corresponding to neuroectoderm/NMPs, mesoderm, and endoderm lineages (**Figure 2F,G**). This organisation reflects the emergence of the three primary germ layers during gastruloid development. Analysis of cell type proportions across conditions revealed a strong effect of Wnt inhibition, with 90% of cells accumulating within a single transcriptional cluster enriched for neuronal signatures (**Supp. Figure S3B**, **Supp. Table 3**). This is consistent with the expected failure to establish a primitive streak–like population due to repression of T expression. In contrast, all germ layer derivatives observed in controls were also present in CDK4/6-inhibited gastruloids (**Figure 2H**), indicating that cell cycle inhibition does not prevent lineage specification. While overall cell type proportions in CDK4/6-inhibited gastruloids remained correlated with controls, measurable shifts in proportion were still present, exceeding twofold for some lineages. These changes included a depletion of mesodermal cells and an increased representation of early progenitor-like states and endodermal cells and ectodermal cells in CDK4/6 condition, confirming prior observations from microscopy and bulk data. Collectively, these findings demonstrate that CDK4/6 inhibition preserves the core differentiation programme but likely perturbs lineage balance, resulting in reproducible changes in germ layer composition. CDK4/6 inhibition appears to affect both cell cycle progression and differentiation processes.

**Table 3:**
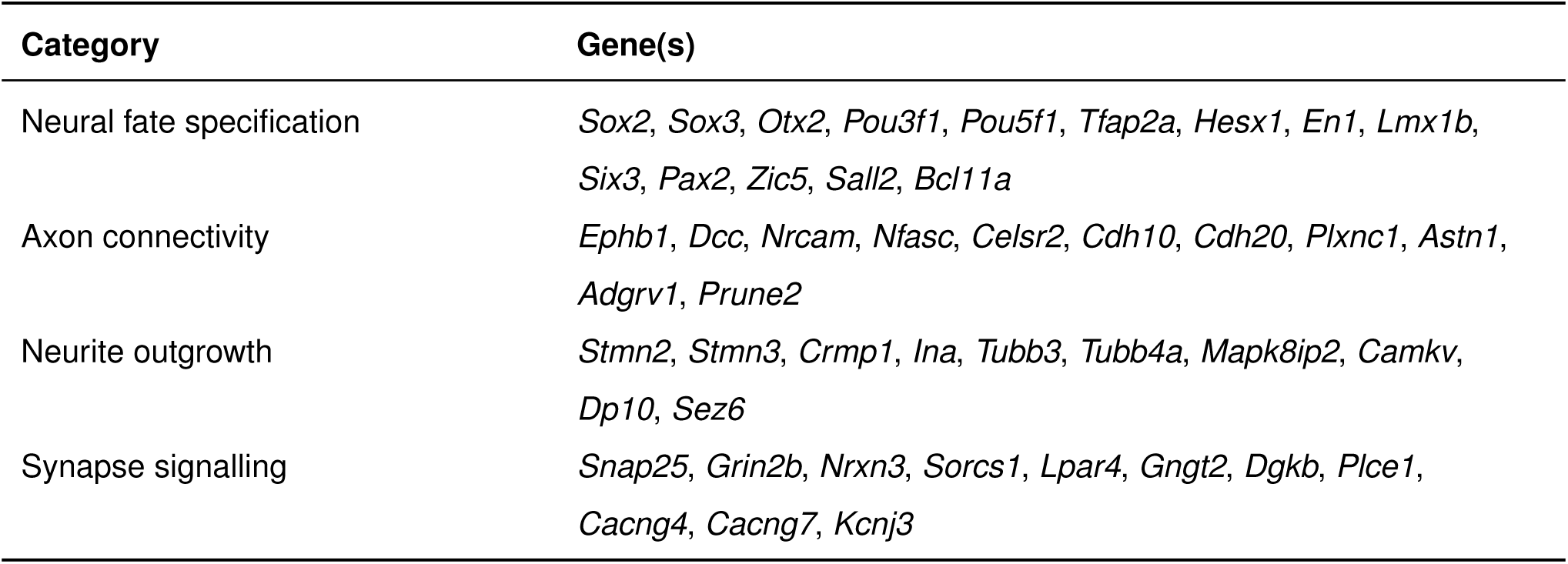
Wnt. inhibition leads to an increased neuronal signature in 120 h gastruloids compared to controls.

However, dimensionality reduction methods like UMAP integrate variation from all sources of cell state and cannot distinguish between the underlying biological processes driving transcriptional changes. Therefore, we sought to disentangle these overlapping sources of variability to better interpret the effects of cell cycle inhibition. To achieve this, we applied a variational autoen-coder (VAE) that separates variance components associated with cell cycle (*θ*-space) from those reflecting other cell states, including differentiation (*z*-space) (**Figure 3A**)^69^. The *θ*-space representation revealed the characteristic circular manifold of the cell cycle, with RNA velocity fields following this cyclic trajectory (**Methods**). This representation further enables inference of a cell cycle phase for each cell. When applied to our dataset, the inferred phase distribution correctly identified the excess of G1-phase cells in the cell cycle inhibited condition (**Supp. Figure S3E**). In contrast, the *z*-space captured the differentiation-related structure of the data, revealing progression of cells from an homogeneous population to distinct branches corresponding to the main germ layers (**Figure 3B**). A closer examination of cell density dynamics across the *z*-space provides further insight into how perturbations affect developmental trajectories (**Figure 3C**). Consistent with previous observations, cells in the control and CDK4/6 inhibition conditions exhibited globally similar temporal progression, whereas Wnt inhibition strongly biased cell fate decisions towards one cell fate. To compare the propensity of cells to follow specific differentiation trajectories, we applied optimal transport analysis (**Methods**), which confirmed that transitions toward mesodermal fates were less frequent under CDK4/6 inhibition (**Figure 3D**), consistent with the reduced mesodermal representation observed by immunostaining and cell-type quantification (**Figure 1F**, **Figure 2H**).

**Figure 3.**
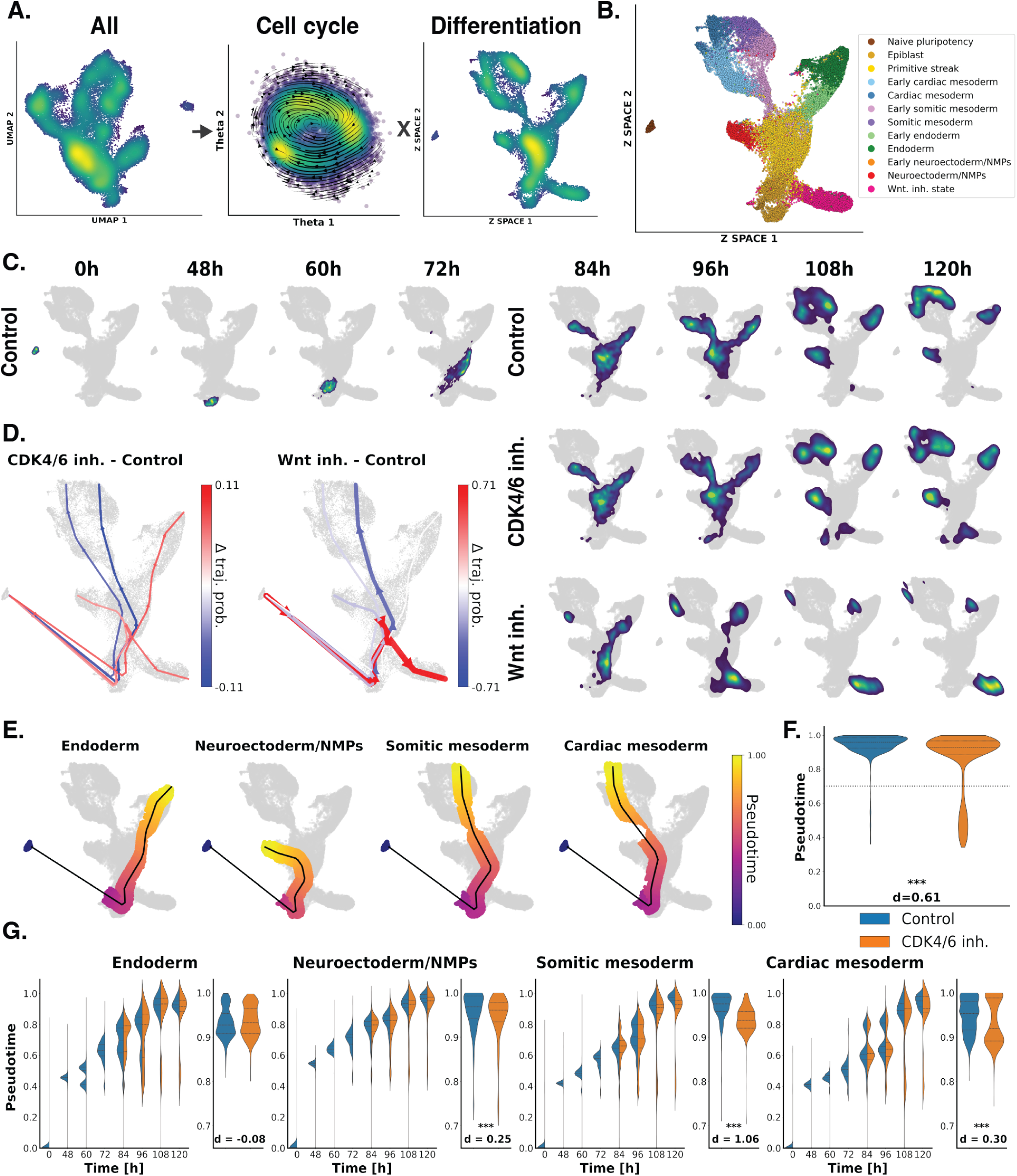
Cell cycle inhibition in gastruloids biases lineage decisions and delays differen-tiation. (A) Disentangled cell cycle (*θ*-space) and cell differentiation (*z*-space) states in the scRNA-seq data, using a variational autoencoder. (B) UMAP projection of the *z*-space shows the differentiation of cells in endoderm, mesoderm, and neuroectoderm/NMPs branches. (C) Cell density, stratified by time and condition, reveals dynamic changes in cell states during gastruloid development. (D) Comparison of trajectory propensities using optimal transport reveals a biased cell fate deci-sion in CDK4/6 inhibitor-treated lineages, characterized by reduced mesoderm differentiation. (E) Trajectories and their associated pseudotime describe progression along different develop-mental paths. (F) Distribution of pseudotime across all lineage trajectories for Control and CDK4/6 inh. condi-tions at 120h highlights the presence of a less differentiated cell cluster (pseudotime *<* 0.7) in the CDK4/6 inh. treated group. Significance levels are indicated as follows: * *p <* 0.05, ** *p <* 0.01, *** *p <* 0.001. ‘d’ denotes Cohen’s effect size. (G) Distribution of pseudotime across different cell type trajectories over time (left) and the asso-ciated pseudotime distribution for 120h gastruloids with pseudotime *>* 0.7 (right) reveal a delay in differentiation, particularly strong in the mesoderm lineages. Significance levels are indicated as follows: *p <* 0.05, *p <* 0.01, *p <* 0.001. ‘d’ denotes Cohen’s effect size.

To further assess differences in the temporal dynamics of differentiation between the control and CDK4/6 inhibition conditions, pseudotime inference was performed across all trajectories, with higher pseudotime values indicating a more differentiated state (**Figure 3E**). Comparing pseudotime distributions between conditions, we observed that in CDK4/6-inhibited gastruloids, a distinct population of cells with very low pseudotime values emerged, corresponding to an epiblast-like state absent in controls (**Figure 3F**). This suggests that some cells failed to commit to a lineage or perhaps reverted to a pluripotent state after cell cycle inhibition.

A second effect we observed was that, even within differentiating populations, the pseudotime distributions of CDK4/6-inhibited cells were globally shifted toward lower values compared with controls, indicating delayed differentiation across lineages. This effect was most pronounced in mesodermal cells but was also evident in neuroectodermal trajectories (**Figure 3G**). However, endodermal differentiation was not significantly affected by this delay.

Overall, scRNA-seq confirmed that gastruloids exposed to CDK4/6 inhibition retain the ability to generate all three germ layer lineages, consistent with immunostaining and bulk transcriptomic data. However, lineage balance is altered, with mesodermal cells consistently underrepresented. Moreover, the presence of a persistent undifferentiated population and reduced pseudotime progression indicate delayed or incomplete differentiation under cell cycle inhibition. Together, these findings show that although normal cell cycle progression is not required to initiate germ layer formation, it influences lineage balance and the efficiency of developmental progression in gastruloids.

### 0.3 RNA velocity and population modelling indicate that cell cycle rate differences alone do not account for CDK4/6-induced cell type rebal-ancing

Cell cycle inhibition resulted in a redistribution of cell-type proportions in gastruloids, suggesting an interplay between proliferation and differentiation. Such changes may arise from an interaction between these processes, or more simply from intrinsic differences in proliferation rates among distinct cell types. Cell cycle inhibition may disrupt these differences, thereby preferentially reducing the proportions of the rapidly cycling populations that may rely on proliferation to maintain their numbers.

To disentangle these effects, we sought to estimate how much of the observed differences in cell-type proportions could be explained by variation in proliferation rates in the different lineages. Yet, our understanding of how cell cycle speed varies between cell types during mammalian development is still limited by a lack of quantitative data. To address this gap, we used our previously published simplified inference of cell cycle periods across cell populations from single-cell data^70^ (**Methods**). This method relies on estimating temporal delays between pre-mRNA and mRNA expression profiles of rhythmic genes, using the previously identified cell cycle phase **Figure 4A**). First, we validated the inferred phases by examining UMI counts, which increased along the cell cycle, consistent with the expected cellular growth over one cell cycle (**Figure 4B**). Likewise, histone transcripts exhibited a distinct peak during S phase, consistent with DNA replication. Moreover, early progenitor cells displayed an enrichment in S/G2/M phases and a marked depletion in G1, consistent with their characteristically shorter G1 phase. Using harmonic regression (**Methods**), genes that exhibited rhythmic expression, defined by a sufficient amplitude in both spliced and unspliced counts, were identified. The genes selected on this basis were largely known cell cycle regulators, and their inferred peak phases were consistent with established phase annotations (**Figure 4C**). Consistent positive shifts were observed between unspliced and spliced counts, reflecting the temporal ordering in which unspliced RNA precedes spliced RNA (**Figure 4D**, **Supp. Figure S4A,B**). The distribution of these shifts across different cell groups was used to estimate the relative cell cycle periods (relative to the half-lives of the genes, **Methods**).

**Figure 4.**
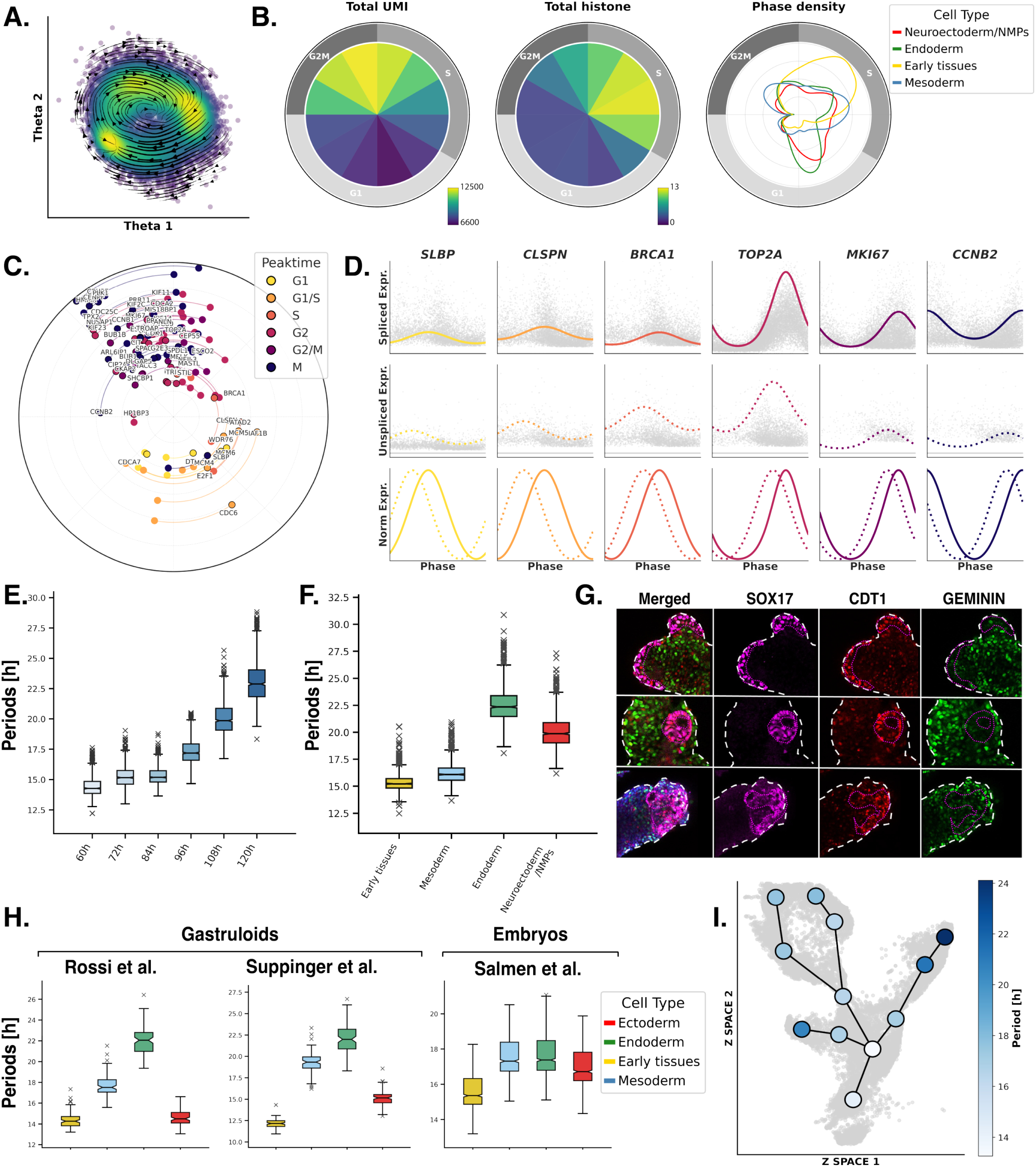
Unspliced-to-spliced RNA shifts allows mapping of cell cycle periods across cell types. (A) The variational autoencoder infers cell cycle phases from scRNA-seq data within a *θ* space, where RNA velocity trajectories are subsequently projected. (B) UMI and histone profiles across cell-cycle phases validate phase assignment, with UMI counts doubling during replication and histone expression peaking in S phase. Phase distri- bution across cell types shows early progenitors are mostly in S and G2M phases. (C) Unspliced and spliced (black outline) transcripts are plotted by their peak phase (angle) and amplitude (radius), linked by arcs; colours indicate annotation of cell-cycle phases from *Cycle-base3* ^77^. (D) Cell-cycle gene profiles reveal systematic phase shifts between unspliced and spliced RNA. (E) Inferred periods show cell cycle lengthening during differentiation. (F) Velocity analysis across cell types reveals longer cycles in endoderm. (G) Fucci gastruloids confirm an overrepresentation of G1 cells in endodermal lineages. (H) Similar cycle length ratios are observed in other public datasets. (I) Finer temporal binning enables mapping of cell-cycle periods along differentiation trajectories.

As an initial validation, this analysis was first applied to different differentiation stages, revealing a progressive lengthening of the cell cycle over time consistent with the expected slowdown of proliferation during differentiation (**Figure 4E**, **Supp. Figure S4C**). Extending the analysis to the cell types revealed distinct proliferation rates across lineages, with endodermal lineages exhibiting slower cycles (**Figure 4F**, **Supp. Figure S4D**). The slower proliferation of endodermal cells was also reflected in an enrichment of G1-phase cells, apparent both in the *θ*-space phase (**Figure 4B, right**) and in Fucci imaging of gastruloids, where Sox17-positive regions were predominantly in G1 (**Figure 4G**). The relative cell cycle periods were further validated using publicly available gastruloid and embryo datasets (**Supp. Figure S4E-I”**). In all cases, the inferred periods increased with differentiation, and the relative proliferation patterns were consistent, with ectoderm cycling faster than mesoderm and endoderm (**Figure 4H**). To characterise how cell cycle dynamics progress along differentiation, period inference was performed at successive nodes along the trajectories, yielding a continuous landscape of generally slowing down proliferations rates along the lineages (**Figure 4I**).

To assess whether the differences in proliferation rates inferred under control conditions could by themselves explain the changes in cell-type proportions observed after CDK4/6 inhibition, we constructed a simple population dynamics model. In this framework, the trajectories were binned into nodes, each representing a segment of the differentiation continuum through which cells progress over time (**Supp. Figure S5A**). The model captures how cell proportions evolve along these nodes by integrating two key processes: proliferation within each node, defined by node *i*-specific growth rates (*g*_i_), and differentiation between node *i* and *j*, described by transition rates (*W*_ij_) (**Figure 5A**). Differentiation transitions were restricted to biologically adjacent nodes, reflecting known lineage progressions (**Figure 5B**). Growth rates (*g*_i_) were fixed to the values inferred from the RNA-velocity model, while the transition rates (*W*_ij_) were fitted to recapitulate the cell proportions observed under control conditions (**Supp. Figure S5B** , **Methods**). This provided a baseline model capturing normal proliferation–differentiation dynamics. To assess the impact of cell cycle perturbation, a second model was simulated to represent the extreme scenario in which proliferation was completely halted from the time of treatment (72h). In this second model, the transition rates were kept constant and taken from the baseline model, while all growth rate values were set to zero from 72h onward, allowing quantification of changes in cell proportions driven solely by the absence of proliferation, independent of changes in the differentiation rates.

**Figure 5.**
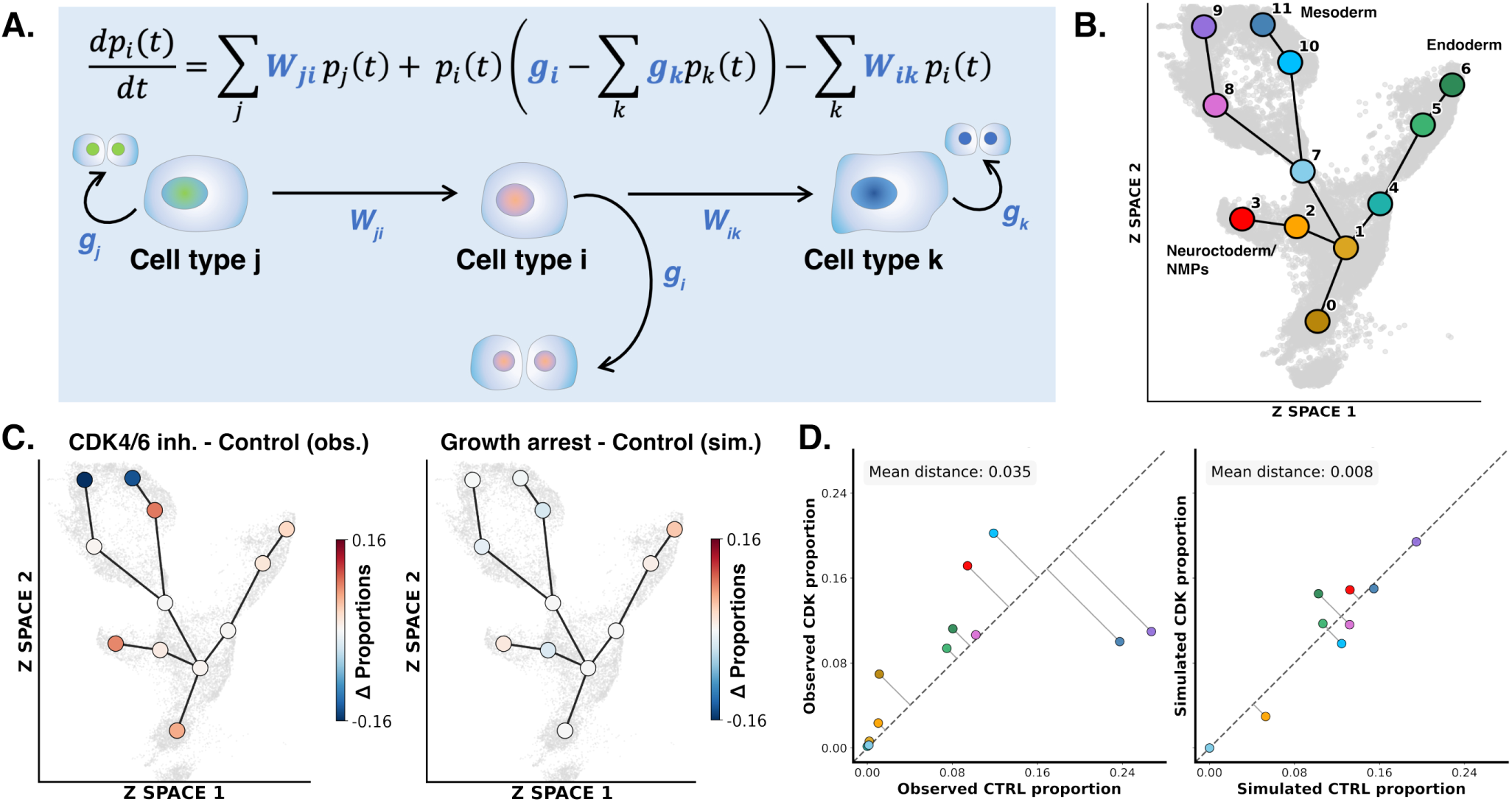
Population modelling suggests that cell cycle rate differences cannot be the sole cause of CDK4/6-induced cell type rebalance. (A) Schematic of the population dynamics model with growth (*g*_i_) and differentiation (*W*_ij_) rates governing cell type proportions. (B) Cell clusters were defined as nodes along the lineage trajectories of the Z-space, for which population densities were modelled over time. (C) Comparison of observed and simulated cell type shifts. CDK4/6 inhibition and growth arrest simulations show changes in the same direction, but CDK4/6 inhibition causes much larger shifts. (D) Scatter plot showing that deviations from the *x* = *y* line are approximately fourfold larger for CDK4/6 inhibition than for growth arrest simulations.

The growth arrest simulation predicted only minor changes in cell-type proportions, indicating that the cell proportions are intrinsically robust to the loss of proliferation alone (**Figure 5C,D**). When compared with experimental data, both simulated and observed trends pointed in the same direction. Slow-cycling lineages such as endoderm were enriched, while fast-cycling tissues like mesoderm were reduced in proportion, consistent with our experimental observations under CDK4/6 inhibition. However, the magnitude of the changes in proportions under CDK4/6 inhibition was approximately fourfold greater than expected from growth arrest alone (**Figure 5D**). Furthermore, the persistence of undifferentiated populations and the delayed progression of differentiation observed in the experimental data of treated gastruloids could not be explained by proliferation differences. Together, these results suggest that intrinsic, lineage-specific cell cycle dynamics account for part of the observed shifts, but additional cell-cycle-dependent mechanisms also alter specific differentiation rates and thus bias cell fate decisions, collectively shaping the full phenotypes seen in CDK4/6-inhibited gastruloids.

## DISCUSSION

Here, we used mouse gastruloids to uncover the developmental consequences of cell cycle inhibition. We found that, despite near-complete growth arrest, key morphogenetic events such as symmetry breaking, elongation, and germ layer patterning proceed largely unperturbed, demonstrating a remarkable robustness of core developmental programmes to proliferation perturbations. Nevertheless, microscopy, single-cell and whole-gastruloid transcriptomics revealed consistent shifts in cell type proportions, most notably a reduction in mesodermal lineages and signs of delayed differentiation. By inferring proliferation rates from scRNAseq data, we show that these changes are only partly explained by cell type–specific cell cycle kinetics, indicating that cell cycle perturbations also modulate differentiation timing and lineage allocation more directly.

### 0.4 Robustness of early development under cell cycle perturbations

A central finding is that gastruloids exposed to CDK4/6 inhibition, although up to ninefold smaller, remain capable of symmetry breaking, axis formation, elongation, and patterning, underscoring strong developmental robustness. Similar robustness to proliferation arrest has been reported in *Zebrafish* ^46–48^, *C. elegans* ^42^, and *Xenopus* ^43–45^, despite some cell type-specific defects^41^.

However, the consequences of cell cycle inhibition on mammalian development have only been sparsely characterised. Although mouse development appears largely robust to knock-outs of cell cycle regulators^49–51^, this does not imply robustness to changes in cell cycle dynamics; rather, it suggests that cell cycle dynamics themselves are robust to the loss of these regulators. For instance, in CDK4/6 double-knockout embryos, which undergo largely normal organogenesis with appropriate cell type proportions yet die later in development due to defective haematopoiesis, cell cycle dynamics are only minimally affected, as reflected by a modest size reduction and normal BrdU incorporation^71^. Compared with the minimal *in vivo* response, CDK4/6-inhibited gastruloids show a strong cell cycle response—including up to a ninefold size reduction, reduced EdU incorporation, and increased G1 occupancy—providing a unique system to examine how real perturbations of cell cycle dynamics influence early developmental programs, a question previously addressed only in specific lineages^53,54^ or later stages^40^.

### 0.5 Cell cycle dynamics as a regulator of lineage balance

In this study, we also observed a systematic rebalancing of cell type proportions under cell cycle inhibition, confirmed across imaging quantifications, individual gastruloid transcriptomes, and single-cell sequencing data. These shifts in proportions were at approximately at most twofold, so five times less important than those reported in zebrafish embryos under similar experimental designs^48^, suggesting greater robustness of cell type proportions in mouse gastruloids. However, differences in the timing of inhibition and in the rates of both cell cycle progression and differentiation between the species may limit the direct comparability of these studies.

Nevertheless, these differences raise the question of whether the cell cycle acts as a regulatory mechanism to establish specific cell type proportions through lineage-dependent dynamics. This notion is closely related to the classical distinction between mosaic and regulative modes of development. In mosaic development, cell fate is largely predetermined by intrinsic factors^72^, whereas in regulative development, fate depends on positional cues and cell interactions^73^. In the context of cell cycle arrest, the question becomes whether cell types proliferate at specific rates to establish the correct proportions, or whether these proportions are set regulatively, independent of lineage proliferation. In litterature, several studies support a role for lineage-specific proliferation in regulating cell type proportions. In *Zebrafish*, fast-cycling lineages are selectively reduced upon cell cycle arrest^48^. Moreover, a recent study demonstrated the importance of apoptosis-mediated cell competition in the gastruloid system, highlighting the role of overall growth rates in shaping cell population dynamics^66^.

Assessing this effect requires first understanding how cell cycle speeds vary between emerging cell types. Therefore, in this study, we propose a new approach to map cell type-specific cell cycle dynamics along differentiation trajectories (**Figure 4**). This method is inspired by a recent development in RNA velocity that leverages unspliced and spliced RNA counts of rhythmic genes to infer cell cycle periods across different cell populations, and whose accuracy has been validated across multiple datasets^70^. While in this study we further validate the approach using complementary analyses (**Figure 4**, **Figure S4**), we acknowledge that the estimates obtained are only relative to to the mean half-life of the cyclic transcripts.

Using the inferred cell cycle rates, we found that fast-proliferating cell types are indeed the most reduced after cell cycle inhibition, suggesting that cell cycle kinetics may partly contribute to the observed shifts in cell type proportions. However, simulations using a minimal model of cell growth and differentiation showed that even in an extreme scenario where proliferation rate differences between cell types were removed, this factor alone could not explain the proportional rebalancing observed under CDK4/6 inhibition (**Figure 5**). Together, these findings suggest that cell type specific cell cycle tuning affects but does not solely determine cell type proportions differences under an inhibited cell cycle.

### 0.6 Mechanisms of cell cycle and cell fate decision coupling

So what other mechanisms could explain the rebalancing of cell type proportions that we observe? Different mechanisms of cell cycle gating of cell fate decisions have been recently reviewed^41^. For example, several studies have reported direct coupling between cell cycle regulators or cell cycle phase distribution and cell fate commitment. Notably, inhibition of CDK4/6 or CDK2 in simpler 2D cellular models, including through Palbociclib treatment, has been shown to impair mesoderm specification, primarily by reducing T expression^27^. Additional studies have reported that Palbociclib can promote neuronal differentiation^30^. Moreover, CDK4/6 activity has been directly shown to suppress endoderm differentiation via the Smad2/Smad3 pathway^34^. Altogether, these findings support a model in which CDK4/6 inhibition favours ectodermal and endodermal fates while reducing mesodermal contributions, which is consistent with our observations. While additional experiments are needed to elucidate the mechanisms involved, the previous study reporting mesodermal impairment had ruled out downregulation of BMP, WNT, or FGF signalling under CDK4/6 inhibition, suggesting that alternative pathways may be at play^27^. One possible mechanism could involve cell cycle regulators–mediated remodelling of chromatin structure, as has been reported previously^25,26,74^.

Another possibility we cannot fully exclude is that the inhibitor exerts off-target effects directly on differentiation regulators, in addition of its specific impact on the cell cycle. However, the bulk sequencing of individual gastruloids performed in this study suggests that cell cycle inhibition with other inhibitors, like the CDK2 inhibitor, leads to a similar rebalancing of cell type proportions (**Figure 1**). Moreover, the study reporting mesodermal impairment under cell cycle arrest also found consistent effects with CDK2 inhibitors and genetic manipulations, including conditional gene knockdowns^27^.

Finally, our findings also highlighted a delayed differentiation under cell cycle–inhibited conditions, characterised by a population that failed to differentiate into any germ layer, as well as a delay within each germ layer, particularly in the mesoderm lineage. This observation contrasts with studies in pluripotent stem cells reporting that inhibition of cell cycle regulators can promote differentiation, suggesting an inverse relationship between proliferation and lineage commitment timing^18–23^. *In vivo*, premature G1/S restriction point has similarly been shown to trigger early neuronal differentiation and disrupt neural tube formation^53,54^. In contrast, other studies have reported effects consistent with our observations, where cell cycle inhibition led to reduced differentiation efficiency, notably within mesodermal derivatives^27^. A global differentiation delay has also been observed *in vivo* at the whole-embryo level in *Zebrafish*. Together, these findings support a context-dependent role of cell cycle regulation in controlling differentiation timing. A possible mechanism underlying the delayed differentiation aligns with some previously proposed models^34^, in which cells must reach a permissive phase of the cell cycle to initiate differentia-tion. A globally slowed cycle could hinder progression to this phase in a fraction of cells, thereby delaying differentiation onset.

### 0.7 Developmental scaling with gastruloid size reduction

Independently of coupling between cell cycle and cell fate decision, the observation that the resulting cell cycle inhibited gastruloids are considerably smaller than control ones raises a broader question of scalability, that is, how the system preserves proportionality in geometric properties, cell type composition, and patterning despite variations in overall size. Such scaling is far from trivial, as gastruloid development is governed by processes such as cell sorting and tissue rearrangements which are also inherently linked to the total number of cells within the system. This phenomenon has been investigated across species^75^ and, more recently, explored in gastruloid models^65,76^, where varying the initial cell seeding number produced gastruloids spanning a wide size range. Although the authors observed systematic, size-dependent timing differences of spatial events, such as delayed symmetry breaking in larger gastruloids, the overall transcriptomic profiles remained robust across sizes. These findings suggest that the transcriptomic differences observed in our case cannot be attributed to size variation, independent of cell cycle effects. Furthermore, the absence of detectable delays in symmetry breaking in our system may reflect the fact that the pronounced size differences emerge only after this developmental event has already occurred.

In summary, this study provides new insights into how early mammalian development responds to cell cycle perturbations. By combining imaging, transcriptomics, and modelling, we show that key developmental processes remain largely robust to near growth arrest, while lineage balance and differentiation timing are systematically altered across data modalities. Our work also introduces an approach to infer cell type–specific proliferation dynamics and highlights other potential interplay between cell cycle regulation and cell fate commitment. Together, these findings refine our understanding of the interplay between proliferation and development of self-organising mammalian systems.

## RESOURCE AVAILABILITY

### Lead contact

Requests for further information and resources should be directed to and will be fulfilled by the lead contact, Felix Naef.

### Materials availability

This study did not generate new materials.

### Data and code availability

- All single-cell and bulk RNA sequencing data has been deposited in GEO under accession number GSE311169 and GSE312366 respectively.
- The code will be provided on a public Github repository upon acceptance.

## ACKNOWLEDGMENTS

We thank members of the Naef group for discussions, particularly Cédric Gobet for his feedback on the manuscript and his help in publishing the transcriptomics data. The CGR8 and Sox1/T reporter mouse ES cell line were kindly provided by the Suter lab in EPFL. We thank the Gene Expression Core Facility, the BioImaging and Optics Platform and the Flow Cytometry Core Facility of EPFL for their support. This work was funded by the Polytechnic School of Lausanne (EPFL).

## AUTHOR CONTRIBUTIONS

M.L. and F.N. conceived the project; M.L. designed, planned and performed experiments. M.L. performed the computational analyses; Y.P. developed the variational auto-encoder for cell cycle phase assignment; M.L. wrote the manuscript with input from all authors. F.N. supervised the project.

## DECLARATION OF GENERATIVE AI AND AI-ASSISTED TECH-NOLOGIES

During the preparation of this work, the authors used ChatGPT-5 to improve clarity, coherence, and linguistic quality of the text. After using this tool, the authors reviewed and edited the content as needed and take full responsibility for the content of the publication.

## STAR METHODS

### Key resources table

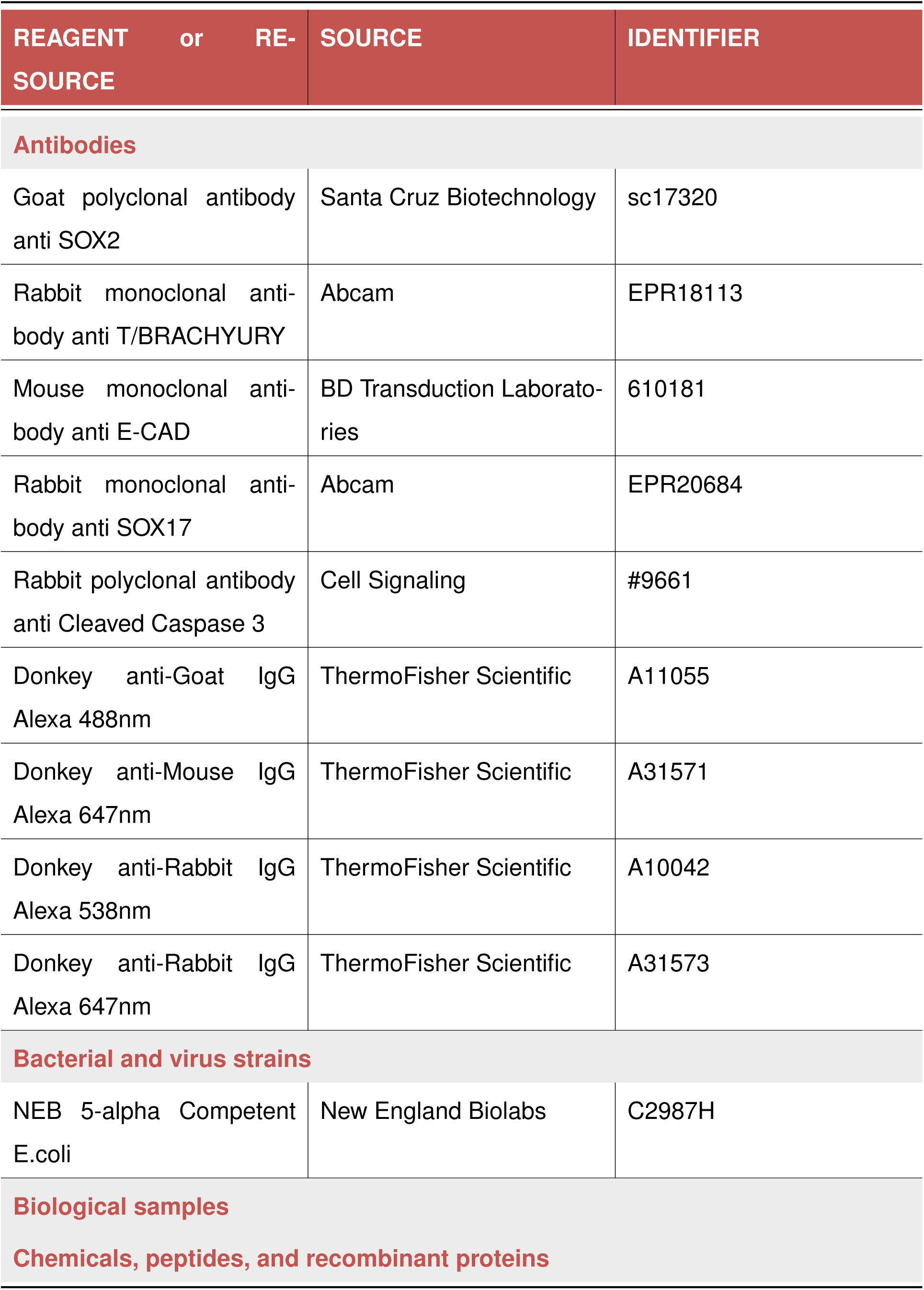
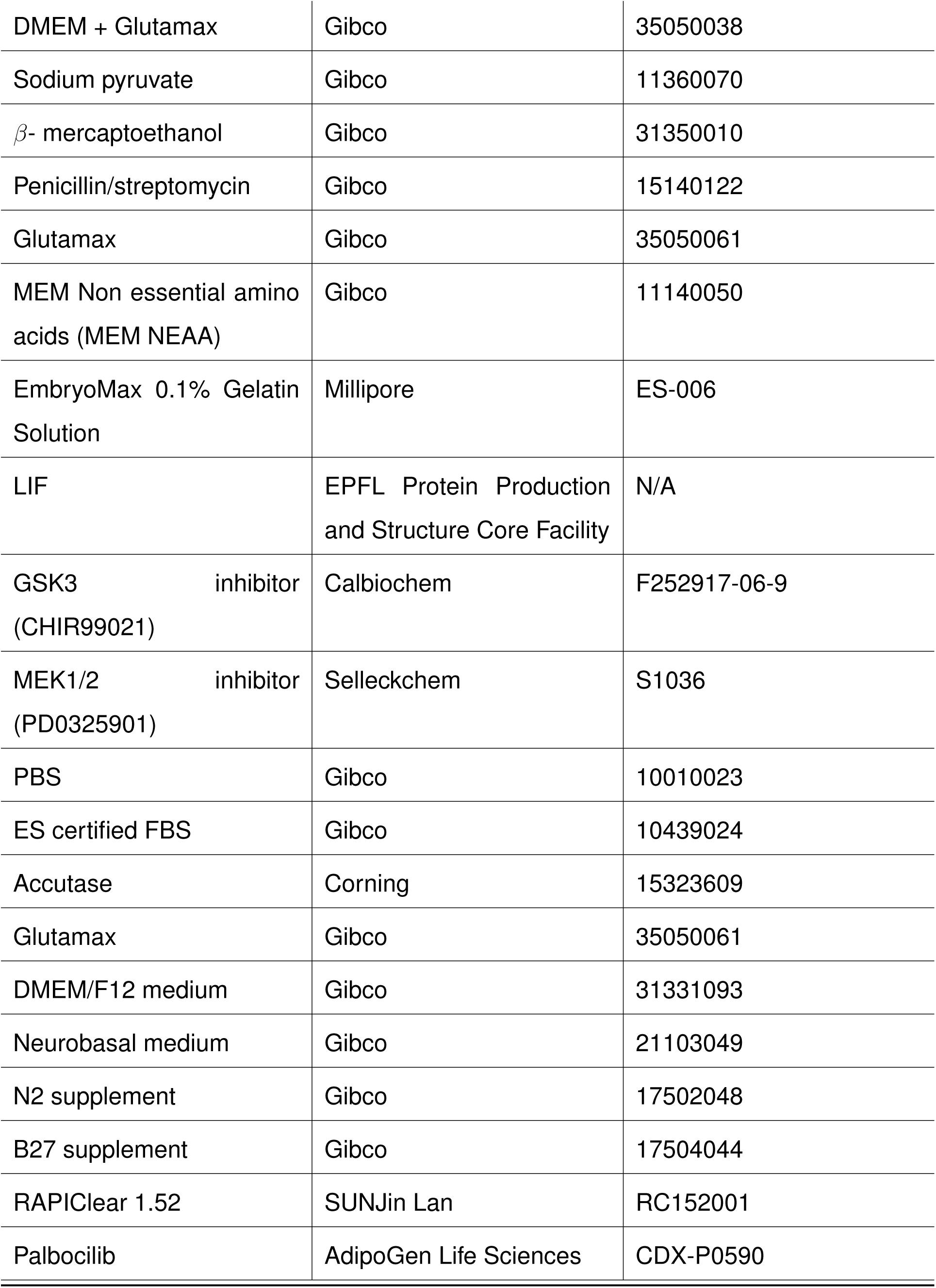
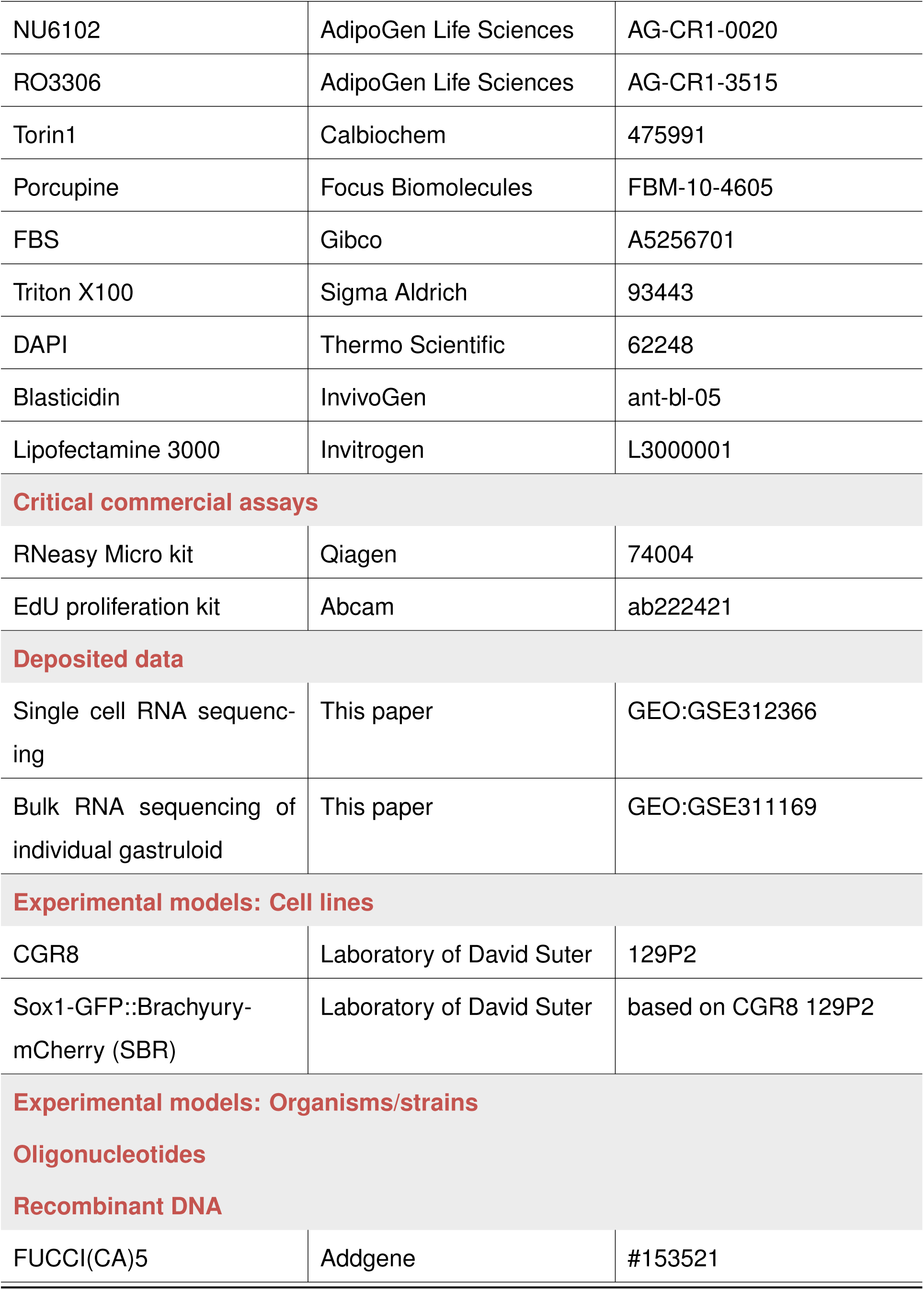
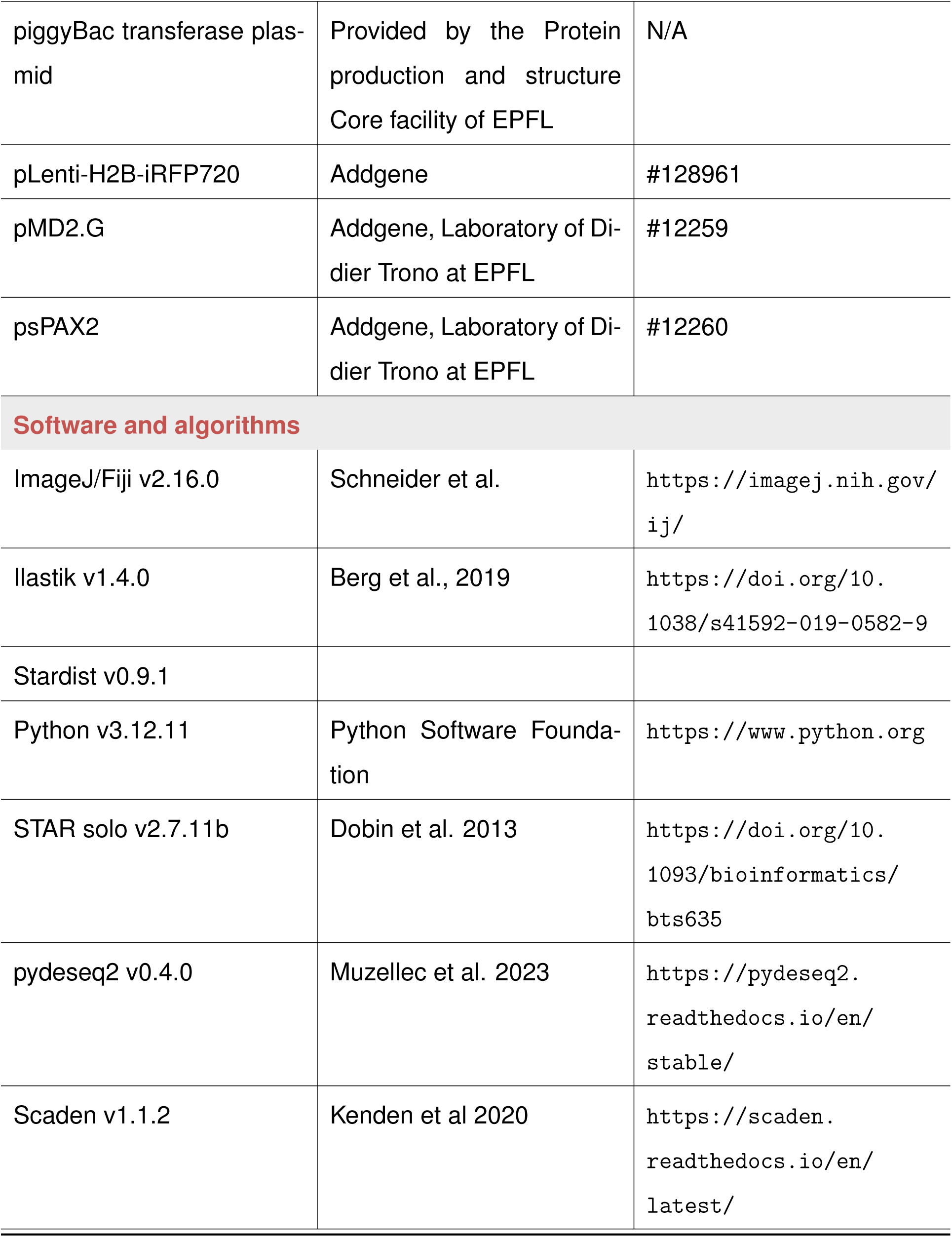
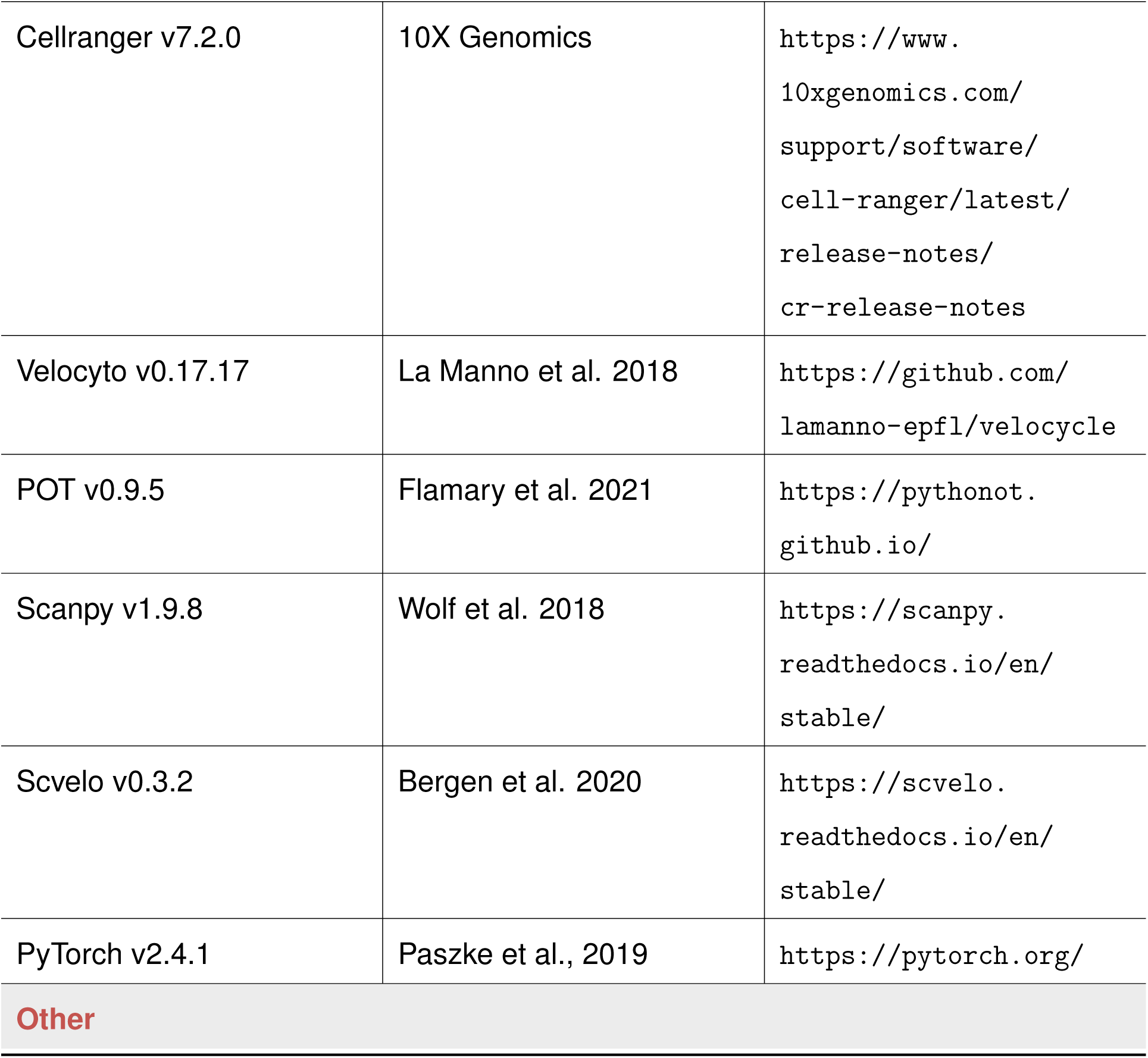

### Method details

#### Cell culture

CGR8 mouse embryonic stem cells were maintained at 37°C in DMEM (*Gibco*, 61965059) supplemented with GlutaMAX (1×; *Gibco*, 35050038), MEM non-essential amino acids (1×; *Gibco*, 11140050), sodium pyruvate (1 mM; *Gibco*, 11360070), penicillin–streptomycin (1×; *Gibco*, 15140122), 2-mercaptoethanol (0.1 mM; *Gibco*, 31350010), and ES-qualified fetal bovine serum (10%; *Gibco*, 10439024). To maintain pluripotency, leukemia inhibitory factor (100 ng/mL; produced by the *EPFL* facility), GSK3 inhibitor (3 µM; *Calbiochem*, 252917-06-9), and MEK inhibitor (PD0325901; 1 µM; *Selleckchem*, S1036) were added to the culture medium. Cells were plated on gelatin-coated plates. Cells were passaged regularly to prevent confluency and spontaneous differentiation.

#### Establishment of cell reporter cell lines

CGR8 embryonic stem cells expressing Sox1-GFP/T-Bra-mCherry (SBR reporter line) were kindly provided by Cédric Deluz from David Suter’s laboratory. To generate the Fucci-expressing line, CGR8 ES cells were transfected with the Fucci(CA)5 (*Addgene*: #153521) plasmid using the PiggyBac transposon system, as previously described^80^. The Fucci constructs and the PiggyBac transposase were co-transfected at a 3:1 ratio using *Lipofectamine*. Transfected cells were selected with *blasticidin* (7 days), and individual -positive single-cell clones were subsequently isolated by fluorescence-activated cell sorting (FACS). In addition, a nuclear reporter was introduced using lentiviral transduction. Lentiviral vectors containing the H2B-iRFP fusion sequence (*Addgene*: #128961) were used in a second-generation lentiviral system. *HEK293T* cells were co-transfected with the lentiviral vector carrying the gene of interest and two packaging plasmids (pMD2.G and psPAX2, obtained from the *Trono* lab), using *HEPES* and *HeBS2X* medium. Viral particles were collected and concentrated 48 h after transfection. CGR8 ES cells were then infected with the produced lentivirus, and H2B-iRFP-positive clones were isolated by FACS.

#### Gastruloid generation

To generate gastruloids, we followed the original protocol described by Van den Brink et al.^59^. ES cells were resuspended in N2B27 medium composed of 50% DMEM/F12 (*Gibco*, 31331093) and 50% Neurobasal medium (*Gibco*, 21103049), supplemented with GlutaMAX (1×; *Gibco*, 35050061), MEM non-essential amino acids (1×; *Gibco*, 11140050), sodium pyruvate (1×; *Gibco*, 11360070), penicillin-streptomycin (1×; *Gibco*, 15140122), 2-mercaptoethanol (0.1 mM; *Gibco*, 31350010), N2 supplement (1×; *Gibco*, 17502048), and B27 supplement (1×; *Gibco*, 17504044). Cells were seeded into 96-well round-bottom plates at a density of 300 cells per well in 40µL of medium. After 48h of incubation at 37°C, a 24h GSK3 inhibitor pulse (*Calbiochem*, 252917-06-9) was applied by adding 150 µL of N2B27 medium containing GSK3i (final concentration, 1×) to each well. After 24h (72h post-seeding), the medium was replaced with fresh N2B27, which was subsequently renewed every 24h.

#### Inhibitors

In this work, we used the various cell cycle and differentiation inhibitors listed below. For the CDK4/6 inhibitor, unless otherwise stated in the manuscript, it was applied at a concentration of 5 µM, starting 72h after aggregation.

| Inhibitor name | Inhibitor target | Reference |
| --- | --- | --- |
| Palbociclib | CDK4/6 inhibitor | *AdipoGen Life Sciences*, CDX-P0590 |
| NU6102 | CDK2 inhibitor | *AdipoGen Life Sciences*, AG-CR1-0020 |
| RO3306 | CDK1 inhibitor | *AdipoGen Life Sciences*, AG-CR1-3515 |
| Torin1 | MTOR inhibitor | *Calbiochem*, 475991 |
| Porcupine | Wnt inhibitor | *Focus Biomolecules*, FBM-10-4605 |

#### Immunofluorescence (IF)

For immuno-fluorescence assays, gastruloids were collected fixed in PFA 4% for 1 hour at room temperature (RT). Gastruloids were then stored in PBS 1X at 4°C. The immunofluorescences were done as follows: After gastruloids were washed 3x in PBS and 3x in PBFST (PBS 1X, 10% FBS, Triton X100 (0.5%)), each for 10min at RT, nonspecific binding of the primary antibody was prevented by a 1-hour blocking step in PBSFT at 4°C. Subsequently, samples were incubated overnight at 4°C in the primary antibodies solution. The different antibodies used, their species and concentrations are below. Gastruloids were washed several times at 4°C in PBSFT (2x 5 min, 3x 15min and 4x 1 hour) and incubated overnight at 4°C in a PBFST solution with DAPI (*Invitrogen*^TM^ 62248, 1:500) and the associated secondary antibodies. Gastruloids were then washed again in PBFST (3x 20min), and then cleared using the RapiClear 1.52 clearing solution (*SUNjin lab*, #RC152002).

Primary antibodies
| Antibody target | Species | Reference | Concentration |
| --- | --- | --- | --- |
| Sox2 | Goat polyclonal | *Santa Cruz Biotechnology*, sc17320 | 1:200 |
| T-Bra | Rabbit monoclonal | *Abcam*, EPR18113 | 1:200 |
| ECad | Mouse monoclonal | *BD Transduction Laboratories*, 610181 | 1:100 |
| Sox17 | Rabbit monoclonal | *Abcam*, EPR20684 | 1:200 |
| Cleaved caspase 3 | Rabbit polyclonal | *Cell Signaling*, #9661 | 1:400 |

Secondary antibodies
| Antibody target | Species | Fluorophore | Reference | Concentration |
| --- | --- | --- | --- | --- |
| Mouse | Donkey | Alexa 647 | *ThermoFisher Scientific*, A31571 | 1:500 |
| Rabbit | Donkey | Alexa 538 | *ThermoFisher Scientific*, A10042 | 1:500 |
| Goat | Donkey | Alexa 488 | *ThermoFisher Scientific*, A11055 | 1:500 |
| Rabbit | Donkey | Alexa 647 | *ThermoFisher Scientific*, A31573 | 1:500 |

#### EdU assays

The EdU assay was performed using the Proliferation Kit (*Abcam*, ab222421) according to the manufacturer’s instructions. At 118 hours after aggregation, half of the medium of the gastruloids (75 µL per well) in control and CDK conditions was replaced with fresh medium (with or without the inhibitor) containing 20 µM EdU, resulting in a final concentration of 10 µM EdU in each well. Gastruloids were incubated with EdU for 2 hours until the 120h endpoint. For microscopy, gastruloids were fixed with 4% PFA for 1 hour. For flow cytometry analysis, gastruloids were collected, washed twice with PBS, and dissociated with *Accutase* for 5 minutes at 37°C. Cells were harvested, washed twice with washing buffer, and fixed in 4% PFA for 20 minutes.

Both fixed gastruloids and dissociated cells were washed twice with a washing buffer and permeabilized using the permeabilization buffer (30 minutes for single cells, 1 hour for gastruloids). For dissociated single-cell samples, each centrifugation step was performed at 300 × g for 5 minutes. Cells and gastruloids were then washed twice, incubated with the reaction mix for EdU detection as described in the manufacturer’s protocol, and stained with DAPI (1:500). Incubation was carried out for 30 minutes for single cells and 1 hour for gastruloids. After two final washes, cells were analysed by flow cytometry, and gastruloids were cleared using the RapiClear 1.52 clearing solution (*SUNjin Lab*, #RC152002) before imaging.

#### Flow cytometry

Flow cytometry measurements (EdU and FUCCI assays) were performed on a *BD LSR-Fortessa*^TM^ equipped with four lasers (405 nm, 488 nm, 561 nm, and 640 nm). Instrument settings were optimised using unstained controls. Cells were manually gated to exclude debris based on forward and side scatter density plots, and positivity thresholds were determined from fluorescence intensity histograms and negative controls.

#### Bulk BRB-seq library preparation of individual gastruloids

Gastruloids were grown from CGR8 cells according to the previously described protocol^59^, with drug treatments applied from 72h onwards: Control (DMSO), CDK4/6 inhibitor (D1: 5uM), CDK2 inhibitor (D1: 10uM, D2: 20uM), and mTOR inhibitor (D1: 1uM, D2: 2uM). For each condition, at least four gastruloids were collected at 120h A.A. from the outer wells of the plate. RNA was extracted from single gastruloids using the *Qiagen RNeasy Micro Kit* (cat. no. 74004). Whole gastruloids were lysed in RLT buffer, and the lysate was homogenized using a *QIAshredder* (*Qiagen*, cat. no. 79656). The homogenized lysate was then passed through an RNeasy MinElute spin column to capture RNA. After assessing RNA quality on the TapeStation, RNA libraries were prepared using the *BRB-seq* protocol (*Alithea* ^81^). Libraries were subsequently sequenced using a 10XG run on the *NovaSeq* platform, targeting approximately 400 million reads.

#### scRNA-seq library generation

For single-cell RNA-seq (scRNA-seq) experiments, multiple batches of gastruloids were generated in parallel from the SBR reporter line, as described below. From 72 hours onward, gastruloids were treated with either DMSO (control), *Palbociclib* (5 µM), or the *Porcupine* inhibitor (3 µM). At the collection time point, gastruloids from the external wells of a 96-well plate were harvested (approximately 40 gastruloids per condition). Samples were washed with PBS and dissociated using *Accutase* and pipetting. Following dissociation, cells were resuspended in PBS containing 0.04% BSA.

Each sample was labelled for 5 minutes according to the 10x Genomics Cell Multiplexing protocol (*Chromium Next GEM Single Cell 3’ v3.1 with Cell Multiplexing*). After three washes in PBS with 1% BSA, cells from different conditions were counted and pooled. The pooled suspension was filtered twice using 40 µm cell strainers (*Sigma*, BAH136800040-50EA) to remove cell clusters and debris.

Single-cell suspensions were processed using the *10x Genomics Chromium* system to encapsulate individual cells in Gel Beads-in-Emulsion (GEMs) and barcode polyadenylated mRNA transcripts for downstream library preparation. Libraries were prepared following the manufacturer’s instructions and sequenced on an *Illumina NovaSeq 6000* platform, targeting approximately 800 million reads per run (100-cycle kit).

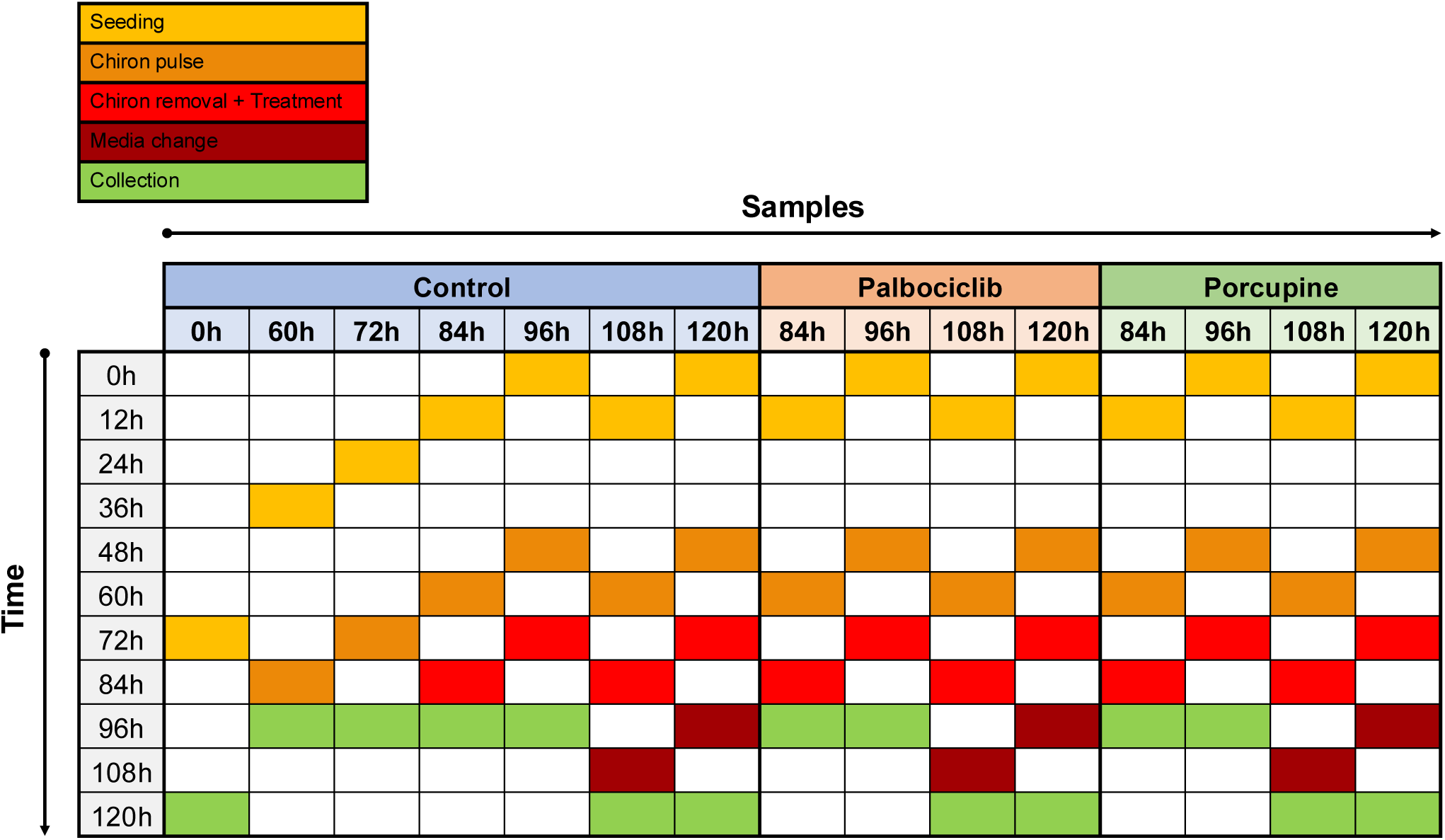

### Quantification and statistical analysis

#### Microscopy and Image quantification

Brightfield live imaging

Brightfield imaging of live gastruloids over time (every 24h) was performed on the *PerkinElmer Operetta* microscope equipped with a 10x/0.35 objective. Gastruloids were segmented using pixel classification in *ilastik* software^82^. Elongation was computed from binary object masks as the ratio of the Feret diameters measured along the major and minor axes, determined by principal component analysis (PCA) of the foreground pixel coordinates.

Fluorescent confocal imaging and quantification

For cell-type marker immunofluorescence, gastruloids, caspase stainings and EdU stainings, gastruloids were imaged using a *Leica SP8* confocal microscope equipped with either a 16x or a 20x immersion objective.

To quantify the distribution of different cell types or caspase levels, each voxel of each z-stack was classified as positive or negative for individual fluorescence channels (DAPI, T, Sox2, and E-cadherin) using intensity thresholding (thresholds were selected manually). Following thresholding, binary masks were generated for each marker and subjected to a median filter to reduce noise. The resulting masks were then combined to assess co-expression patterns between markers. Ratios of different cell populations were calculated as the number of pixels positive for a given marker (or marker combination) divided by the total number of DAPI-positive pixels.

Cell cycle phase quantification in Fucci gastruloids was performed as follows. Cells were segmented independently in each Z-slice using *StarDist* ^83^, first on the sum of the red and green fluorescence channels, and then separately on each channel. A cell was considered positive in the red or green channel if the centroid of the cell identified in the summed image also fell within a segmented region of the corresponding individual channel.

#### RNA-seq data analysis of the individual gastruloids

Raw reads from the BRB-seq experiment were demultiplexed and mapped using *Alithea BRB-SeqTools*. Differential gene expression analysis and CPM-based comparisons between samples were performed using the *DESeq2* library^84^ implemented in Python.

For cell-type deconvolution, the algorithm described in *Scaden* was applied following the authors’ instructions^68^. The *Scaden* model was first trained on our own scRNA-seq dataset (control condition, 120h) using our cell-type annotations, and subsequently applied to the bulk RNA-seq expression matrix to estimate the proportion of each cell type.

#### scRNA-seq data analysis

Raw sequencing reads were mapped to the reference genome using *Cell Ranger* ^85^ (10x Genomics). Spliced and unspliced transcript counts were then extracted and demultiplexed using the *Velocyto* pipeline^86^. Data processing and quality control were carried out using the *Scanpy* ^87^ library in Python. Cells with low total UMI counts or a high proportion of mitochondrial gene expression were filtered out to exclude low-quality or dying cells. Lowly expressed genes were also removed. The counts were then normalised per cell using total count normalisation and log-transformed for subsequent analysis.

For cell type assignment, cells were first clustered using the *Louvain* algorithm, after which cell type markers (list generated from^64^ and^78^ datasets) were used to manually annotate each cluster.

#### Estimation of latent cell state and cell cycle phases

From the scRNA-seq data, we estimate cell cycle phases for each individual cell, while accounting for context-specificity arising from biological factors such as cell type.

For this, we used a variational autoencoder (VAE), with a structured latent space consisting of a cell cycle phase (*θ*), and a cell-context variable *z*, as described in^69^.

Briefly, the model^69^ uses as input a set of 98 cell cycle genes, and the 2000 most variable genes in the dataset. The variable genes are projected onto a ten-dimensional latent coordinate *z*, using the mean and variance of an isotropic multivariate Gaussian distribution. The cell cycle genes project onto a two-dimensional space from which the angle is modelled as a power spherical distribution^88^. For each gene *g*, the latent variables are then decoded into the sum of an aperiodic component and a circular component consisting of a Fourier series.

#### scRNA-seq trajectory reconstruction

To reconstruct temporal trajectories of cells within the *z*-space, we employed optimal transport (OT) to link cell distributions between successive time points for each experimental condition, using the *Python Optimal Transport* ^89^ library. A common set of baseline cells at 0h was randomly selected to initialise representative trajectories. For each condition, cell coordinates at consecutive time points were treated as source and target distributions. Pairwise squared Euclidean distances between all cells at time *t* and time *t*+1 were computed to form the transport cost matrix, and the Earth Mover’s Distance (EMD) solution was then obtained to derive the optimal transport plan that minimises the overall transport cost. This procedure was applied iteratively across all consecutive time intervals, yielding sets of synthetic trajectories that approximate the most probable flow of cell populations through the embedding over time. The resulting trajectories were subsequently projected onto a lower-dimensional representation using principal component analysis (PCA) and clustered with k-means to group similar patterns. Each cluster corresponded to a putative lineage, which we annotated manually. Finally, we quantified and compared the number of trajectories in each cluster across conditions to reveal biases in lineage commitment and cell-state transitions. For each trajectory cluster, we computed the mean trajectory, which served as the reference path for pseudotime assignment. For these mean tra-jectories, we calculated the shortest Euclidean distance from each cell’s position in *z*-space to the piecewise linear curve defined by the trajectory points; cells within a fixed distance threshold were considered associated with that trajectory. For each associated cell, pseudotime represents how far along the trajectory the cell lies, relative to its start and end. To compute this, we measured the total length of the trajectory and the distance along the path from the start to the point on the trajectory closest to the cell (its projection). The pseudotime value is then this distance divided by the total trajectory length, giving a normalised value between 0 (trajectory start) and 1 (trajectory end).

#### Simplified RNA velocity model for cell cycle period inference

We here used our previous simplified RNA velocity model^70^ to estimate cell cycle periods across the lineages, which is based on the delays between mRNA and pre-mRNA accumulation profiles. To characterise those delays genome-wide, we used the cell cycle phases (*θ*_c_) inferred from the VAE and fitted each gene with a harmonic regression (one harmonic) using a generalized linear model (GLM) with Poisson noise. For each gene, expression levels across cells were modelled as a one-harmonic Fourier series, specifically the expected mean counts was taken as

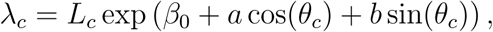

where *L*_c_ denotes the total transcript number per cell (used as an offset) and [*β*_0_*, a, b*] are the gene specific regression coefficients to be estimated. Model parameters were estimated for each gene individually using iteratively re-weighted least squares (IRLS). The model was first applied genome-wide to the spliced transcripts to identify genes with clear periodic mRNA profile. Fitted log-amplitudes greater than 0.2 (log) were selected for further analysis. Subsequently, the same model was refitted on the unspliced transcript layer for the subset of oscillatory genes. Genes with unspliced amplitudes exceeding 0.1 (log) were kept for further analyses, to ensure periodic spliced and unspliced expression.

As in^70^, the phase shifts Δ*ϕ*_sg_ between unspliced and spliced mRNA in condition s then depend on the transcript half-lives *τ*_g_ = *γ*^−1^ and frequencies *ω*_s_ = 2 ∗ *π/T*_s_ (*T*_s_: period) according to

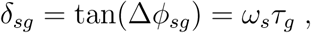

To compare conditions, we selected on genes that were rhythmic in both spliced and un-spliced counts across all conditions *s*.

This defines a rank-1 decomposition of the matrix *δ*_sg_. Using the singular value decomposition (SVD)

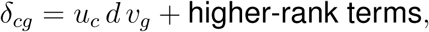

we can obtain condition-specific cell-cycle velocity

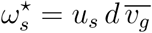

where *v*_g_ denotes the mean over genes. The corresponding cell-cycle period, expressed in units of mean transcript half-lives, is then given by

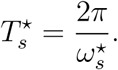

For all analyses, the mean transcript half-life *v*_g_ was fixed to *τ* = 1.25 h, consistent with typical mRNA decay times in mammalian single-cells.

#### Cell type proportion model

Using the above trajectory reconstruction, we coarse-grained the cells into discrete nodes (11 nodes) along the different differentiation trajectories and for each inferred their growth rates using the simplified RNA velocity model. We also modelled biologically plausible transitions between nodes based on established biological knowledge. To describe the temporal evolution of cell proportions across these nodes, we employed a linear ordinary differential equation (ODE) model capturing net cell growth and differentiation dynamics. The number of cells in node *i* at time *t*, denoted *N*_i_(*t*), evolves according to

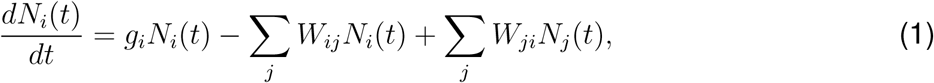

where *g*_i_ represents the *effective growth rate* of node *i*, accounting for the combined effects of proliferation and apoptosis, and *W*_ij_ denotes the differentiation rate from node *i* to node *j*.

Because scRNA-seq data can only be used in relative rather than absolute cell numbers, we reformulated the model in terms of the *proportion* of cells in each node at time *t*:

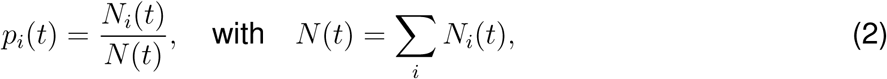

which yields the proportional model

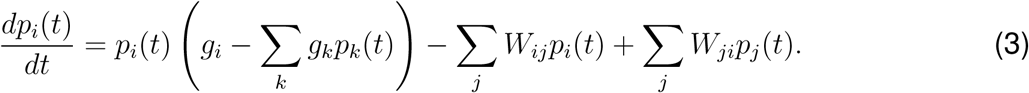

We note that the proportions are invariant with respect to a global shift in the growth rates (*g*_i_ → *g*_i_ − Δ).

We implemented the model in Python, solving the system of ODEs with the scipy package.

The transition matrix (*W* ) was fitted on the control condition data, after which we simulated a sce-nario of complete growth arrest by setting all node-specific growth rates to zero after 72 h while keeping the transition dynamics unchanged. The simulated trajectories were then compared to both the observed and fitted control trajectories to assess the impact of growth suppression on differentiation dynamics.

## SUPPLEMENTARY FIGURE TITLES AND LEGEND

**Figure S1.**
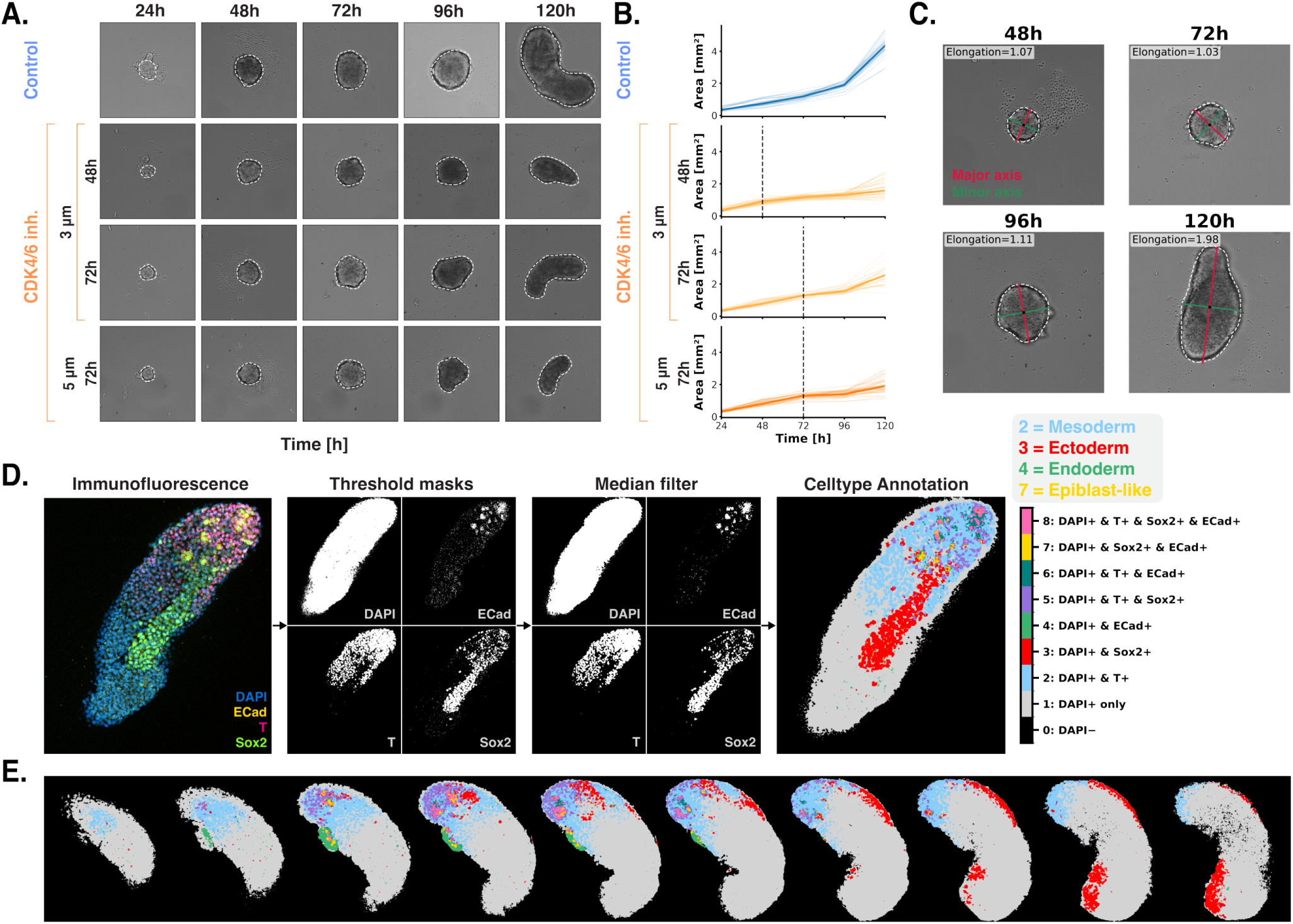
Quantification of gastruloid growth, elongation, and cell-type composition from microscopy images. (A) Brightfield Operetta imaging tracks gastruloid morphology over time; white overlays show segmentation masks used for area quantification. (B) Area progression curves of individual gastruloids over time demonstrate high size repro-ducibility within conditions. *n >* 20 for all conditions. (C) Elongation is quantified as the ratio of the major to minor axis, both fitted to the mask of gastruloid brightfield images. (D) IF staining quantification pipeline. For each image z-stack, per-channel thresholds (DAPI, E-CAD, T, SOX2) classify pixels as positive or negative. A median filter removes noise. Pixels are then labelled by co-positivity: T-only = mesoderm; E-CAD-only = endoderm; SOX2-only = neuroectoderm/NMPs; SOX2+E-Cad = early tissues (epiblast-like). (E) IF staining quantification is performed over the full z-stack for each gastruloid.

**Figure S2.**
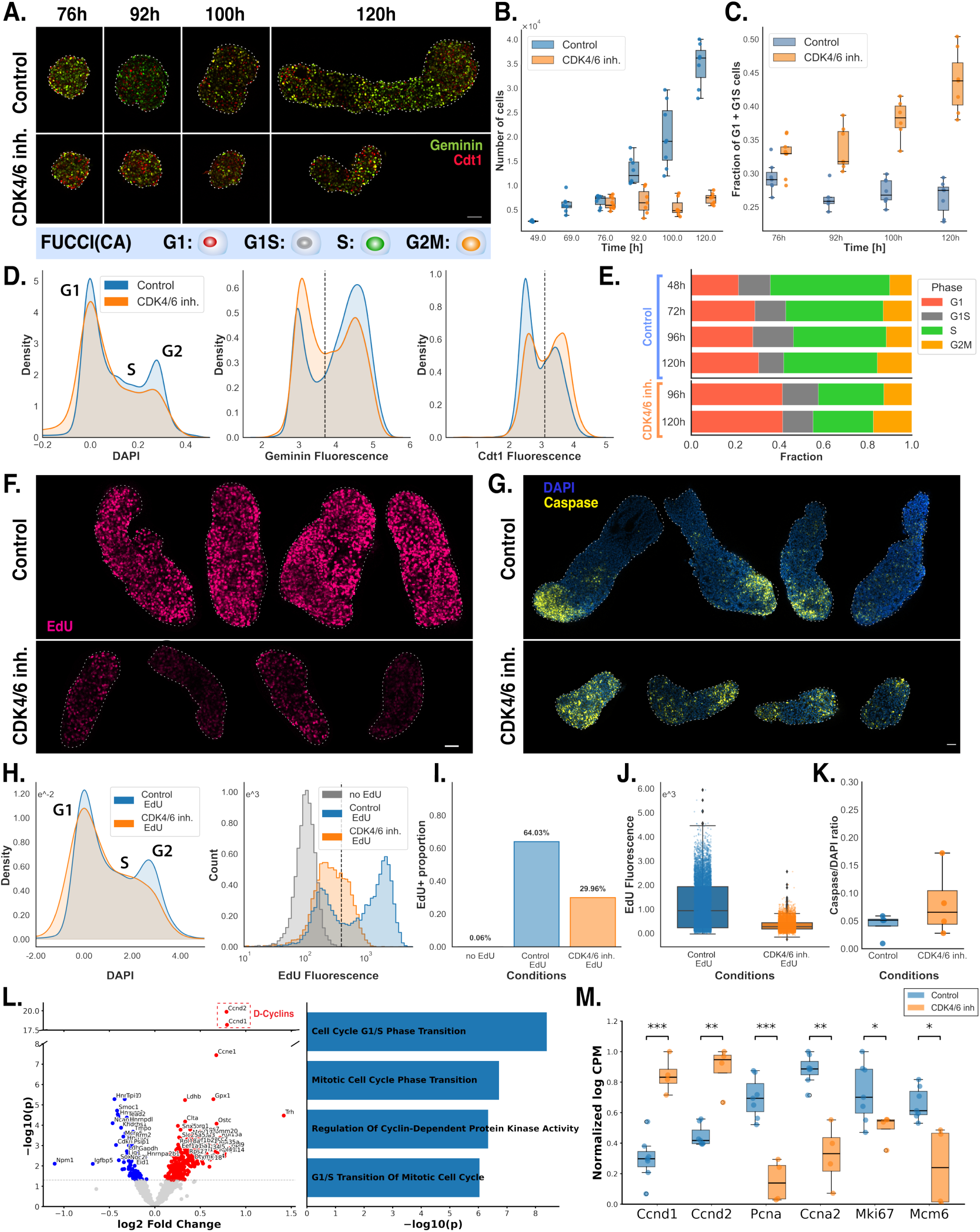
Cell-cycle characterisation of CDK4/6-inhibited gastruloids. (A) Fucci(CA) imaging of gastruloid across time allows characterization of cell-cycle phase dis-tributions in Control and CDK4/6 inh. gastruloid. Red cells are in the G1 phase, green cells in the S phase and orange cells are in the G2M phase. (B) Quantification of cell number from Fucci(CA) gastruloid images shows a growth arrest under 5um CDK4/6 inhibition. (C) Quantification of cell cycle state from Fucci(CA) gastruloid images shows a significant, time-dependent increase in G1/G1–S cell proportions under 5um CDK4/6 inhibition. (D) Flow cytometry fluorescence histograms (DAPI, GEMININ, CDT1) allows characterization of the population cell-cycle state in control and CDK4/6 inhibited gastruloid; black dotted lines indicate positivity thresholds. (E) Cell cycle phase proportions quantified from flow cytometry confirm enrichment of G1 and G1/S cells and reduction of S and G2/M cells in the cell cycle inhibited condition. (F) Representative EdU incorporation imaging in 120h gastruloid reveals reduced incorporation under CDK4/6 inhibition compared with control. (G) Representative Caspase stainings illustrating the identification of apoptotic cells in control and CDK4/6 inh. gastruloid. (H) Flow cytometry fluorescence histograms (DAPI, EdU) identify EdU-positive cell population; black dotted lines indicate positivity thresholds. (I) Quantification of EdU-positive fractions shows a significant decrease under CDK4/6 inhibition compared to the control condition. (J) Distribution of EdU fluorescence in EdU-positive cells shows a shift to lower intensities under CDK4/6 inhibition compared with controls, consistent with slower S-phase progression. (K) Caspase stainings quantification shows a higher, but variable, fraction of dying cells under inhibition. n=4 in each condition. (L) Differential expression and GO term analysis of bulk RNA-seq from individual gastruloids reveal differences in cell-cycle pathways between conditions. _(M)_ Normalised log-CPM of cell-cycle genes shows up-regulation of D-cyclins (G1) and down-regulation of proliferative markers (e.g., *Mki67*, *Pcna*) with cell cycle inhibition. *n*_control_ =7, *n*_others_ = 4.

**Figure S3.**
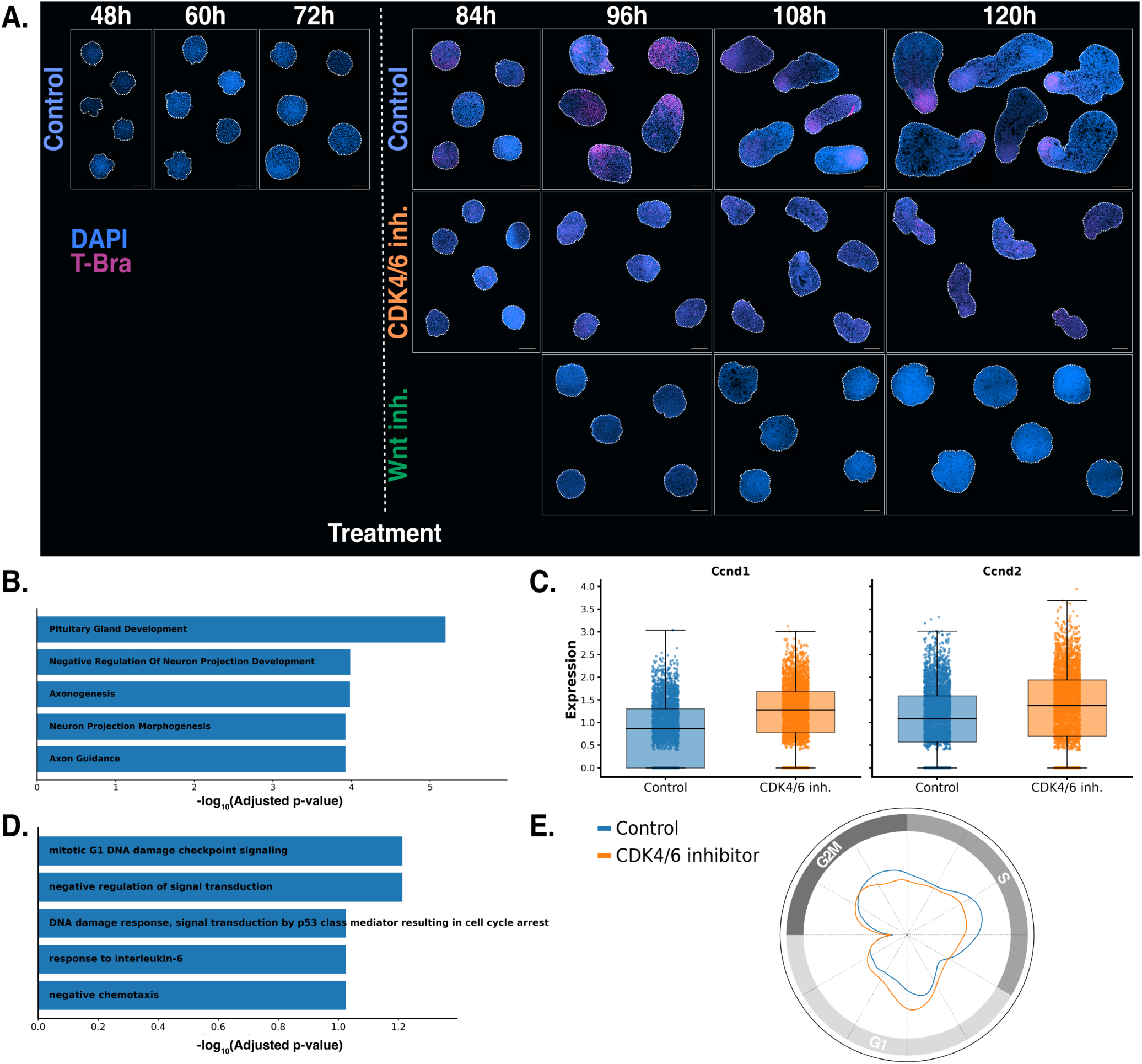
Characterisation of single-cell transcriptomic responses to CDK4/6 and Wnt inhibition in developing gastruloids. (A) Representative T-mCherry–expressing gastruloids from the same batches used for scRNA-seq, shown at the time of collection. (B) Enrichment of neuronal GO terms in the PORCN-inhibited cluster indicates Wnt inhibition promotes neuronal differentiation. (C) D-Cyclin overexpression under CDK4/6 inhibition confirmed by scRNA-seq. (D) GO term analysis of CDK4/6 inhibitors–upregulated genes validates G1 cell-cycle deregula-tion. (E) *θ*-space phase density confirms an excess of G1-phase cells under CDK4/6 inhibition.

**Figure S4.**
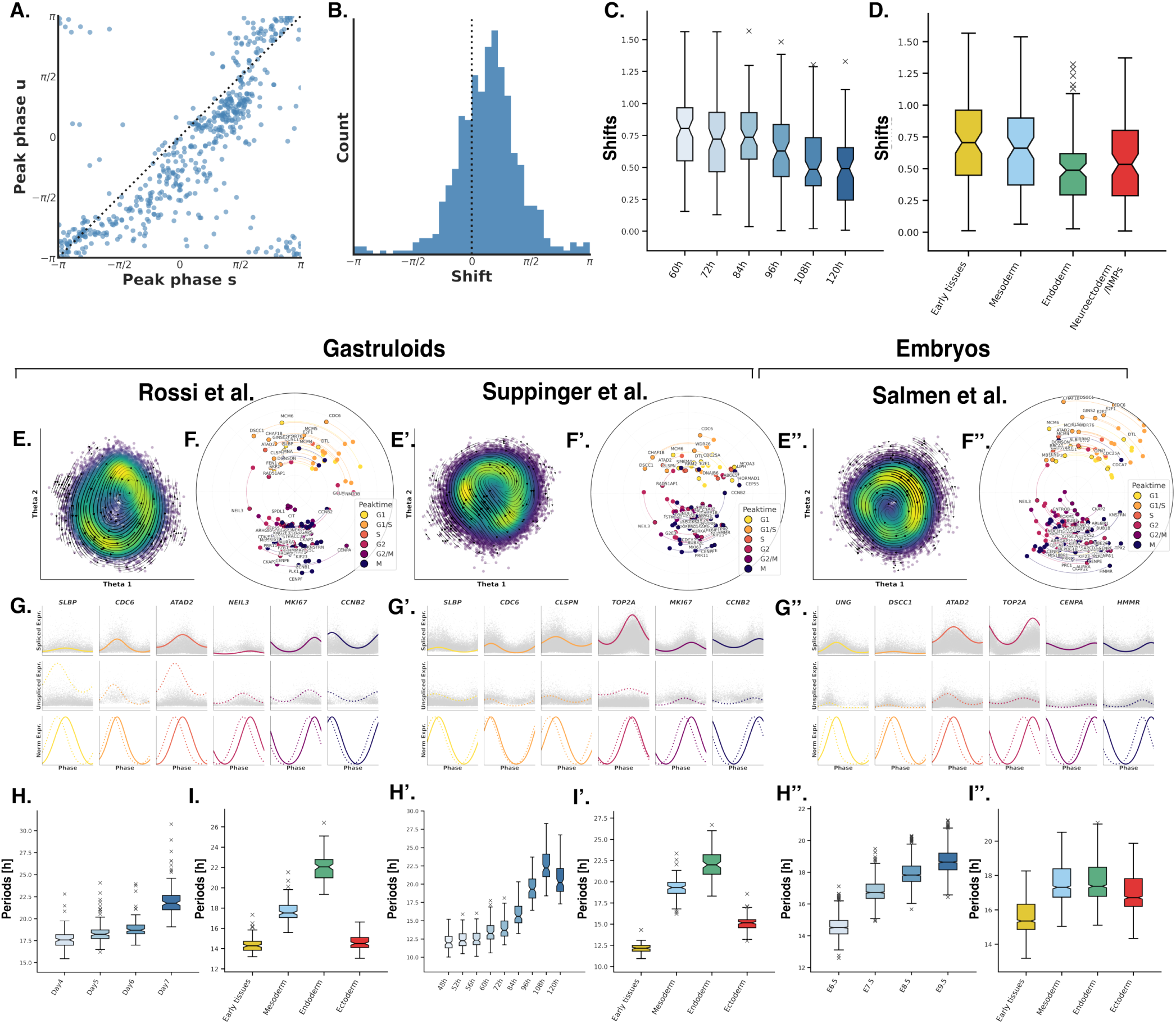
Characterisation and validation of cell cycle period mapping across datasets. (A) Scatter plot of unspliced versus spliced RNA peak times, with most genes falling on one side of the diagonal, indicating consistent phase shifts between the two. (B) Distribution of phase shifts, with most genes exhibiting shifts between 0 and *π/*2. (C) Phase shift distribution over time reveals a negative correlation between phase shift and dif-ferentiation time, consistent with cell cycle lengthening during differentiation. (D) Phase shift distribution across cell types highlights distinct patterns among cell types. For the Rossi et al.^78^, Suppinger *et al.*^64^ (‘), and Salmen et al.^79^ (‘’) datasets: (E) Projection of cells in the inferred circular *θ*-space obtained from the variational autoencoder model and RNA velocities trajectories. (F) Polar plots displaying unspliced and spliced (outlined) transcript profiles by peak phase (an-gle) and amplitude (radius); arcs link corresponding spliced and unspliced counts of the same gene, and colours indicate annotated cell cycle phases. (G) Unspliced and spliced RNA expression profiles of cyclic genes plotted as a function of cell cycle phase. (H, I) Boxplots and scatter plots showing inferred cell cycle period distributions and correspond-ing cell velocities across cell types and lineages.

**Figure S5.**
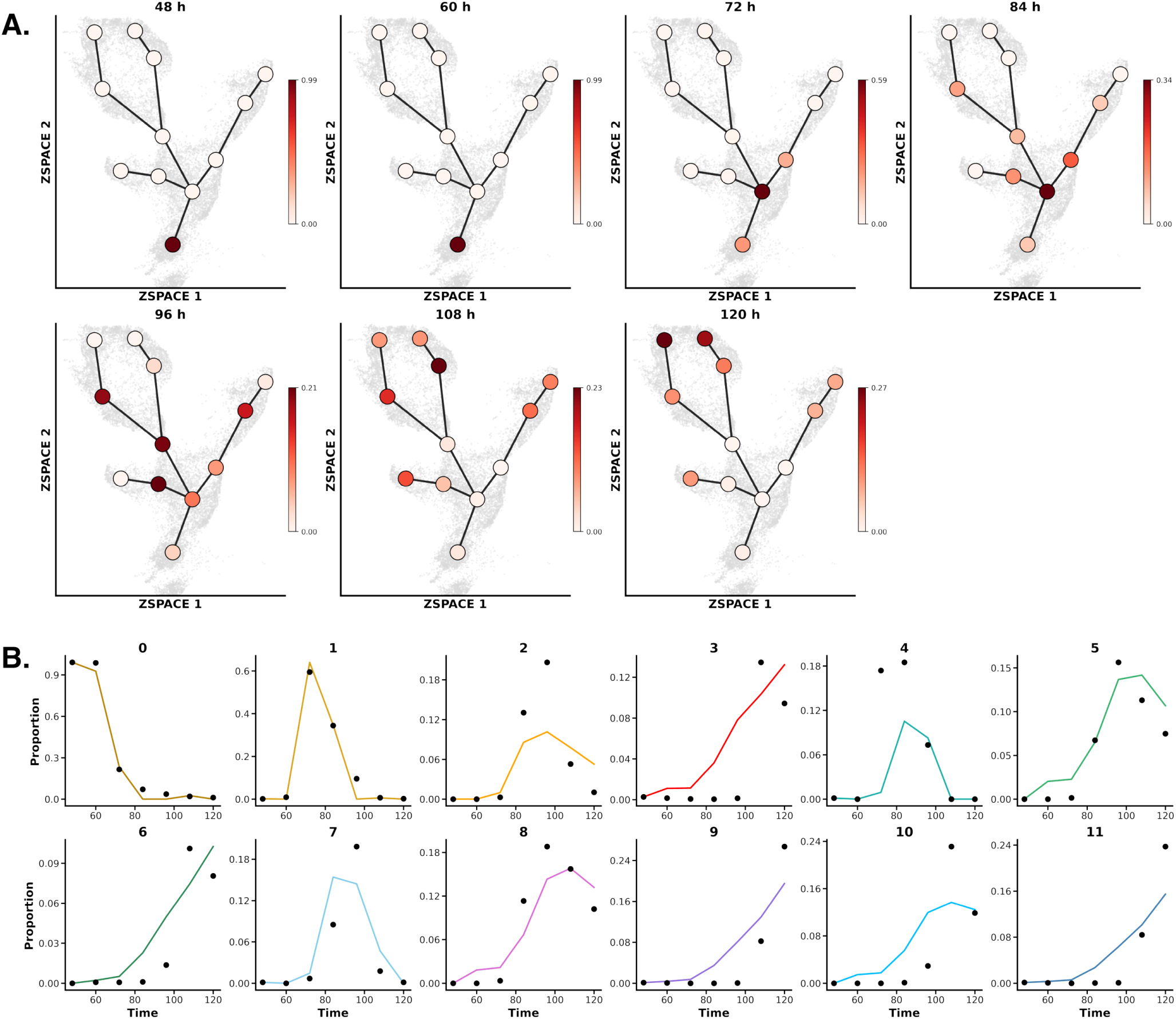
Simulation of Population Dynamics Under Control and Growth Arrest Condi-tions. (B) Temporal evolution of observed population densities across UMAP-defined node compart-ments in the Control condition. (C) Observed control proportions are shown as black dots, with their simulated fits plotted as coloured lines over time. Numbers correspond to nodes illustrated in Figure 5B.

